# Uncovering the Interleukin-12 Pharmacokinetic Desensitization Mechanism and Its Consequences with Mathematical Modeling

**DOI:** 10.1101/2024.02.20.581191

**Authors:** Jonathon DeBonis, Omid Veiseh, Oleg A. Igoshin

**Affiliations:** Department of Bioengineering, Rice University, Houston, Texas, USA; Center for Theoretical Biological Physics, Department of Chemistry, Department of Biosciences, Rice University, Houston, Texas, USA

**Keywords:** interleukin-12, pharmacokinetics, mathematical modeling, cancer immunotherapy, drug transport, dynamical systems analysis

## Abstract

The cytokine interleukin-12 (IL-12) is a potential immunotherapy because of its ability to induce a Th1 immune response. However, success in the clinic has been limited due to a phenomenon called IL-12 desensitization – the trend where repeated exposure to IL-12 leads to reduced IL-12 concentrations (pharmacokinetics) and biological effects (pharmacodynamics). Here, we investigated IL-12 pharmacokinetic desensitization via a modeling approach to (i) validate proposed mechanisms in literature and (ii) develop a mathematical model capable of predicting IL-12 pharmacokinetic desensitization. Two potential causes of IL-12 pharmacokinetic desensitization were identified: increased clearance or reduced bioavailability of IL-12 following repeated doses. Increased IL-12 clearance was previously proposed to occur due to the upregulation of IL-12 receptor on T-cells that causes increased receptor-mediated clearance in the serum. However, our model with this mechanism, the accelerated-clearance model, failed to capture trends in clinical trial data. Alternatively, our novel reduced-bioavailability model assumed that upregulation of IL-12 receptor on T-cells in the lymphatic system leads to IL-12 sequestration, inhibiting the transport to the blood. This model accurately fits IL-12 pharmacokinetic data from three clinical trials, supporting its biological relevance. Using this model, we analyzed the model parameter space to illustrate that IL-12 desensitization occurs over a robust range of parameter values and to identify the conditions required for desensitization. We next simulated local, continuous IL-12 delivery and identified several methods to mitigate systemic IL-12 exposure. Ultimately, our results provide quantitative validation of our proposed mechanism and allow for accurate prediction of IL-12 pharmacokinetics over repeated doses.

## Introduction

Inteleukin-12 (IL-12) is a heterodimeric cytokine consisting of the p35 α subunit and the p40 β subunit^1^. The unmodified heterodimer form of IL-12 has a molecular weight of 58.9 kDa; however, it is glycosylated and migrates as a 65-75 kDa protein in SDS PAGE^2^. IL-12’s immunological activity causes differentiation of CD4+ helper T-cells (Th-cells) into the Th1 phenotype, leading to the production of pro-inflammatory cytokines and immune system activation^3^. The primary cytokine induced by IL-12 activity is IFNγ, but TNFα, GM-CSF, IL-2, and IL-8 are also known to be produced in response to IL-12 stimulus^1,4^. Immune activation is achieved through the binding of IL-12 to its receptor, which ultimately leads to the activation of STAT4^1,4,5^. IL-12’s ability to trigger robust immune system activation motivates its investigation as a cancer immunotherapy.

The first clinical trial investigating IL-12 was initiated in 1997 for the treatment of cancer patients with advanced malignancies^1,6^. Here, patients were first treated with an i.v. test dose of IL-12 two weeks before receiving five times weekly i.v. doses of IL-12 every 28 days^6^. A phase two trial was initiated shortly after following the same dose regimen but without the test dose. In contrast to the phase one, this trial was terminated after severe toxicities, including treatment-related patient deaths^7^. It was apparent that prior exposure to IL-12 desensitizes patients to IL-12 activity, evident through the ∼5-fold difference in measured serum IFNγ in patients treated with the same dose between trials.

IL-12 desensitization has been observed in a large number of additional clinical trials, with both IL-12 pharmacokinetic (PK) and pharmacodynamic (PD) desensitization observed^8–17^. PD desensitization is characterized by reduced IFNγ levels following repeated doses^6,7,13,16^, whereas PK desensitization causes reduced IL-12 concentrations.^14,15,18^. Furthermore, PK and PD desensitization appear to be caused by separate mechanisms due to a difference in the timescale in which these phenomena occur. In a study of repeated subcutaneous IL-12 doses in mice, measured serum IL-12 concentrations decreased with every dose administered, whereas measured serum IFNγ concentrations decreased after three doses were administered^18^. Despite increasing tolerance to IL-12 administration, studies have shown that IL-12 efficacy is also hindered by desensitization^11,13,19^.

Several mechanisms have been proposed in the literature to explain IL-12 PK and PD desensitization; however, none have been validated quantitatively. Importantly, evidence in literature suggests that IL-12 desensitization is not due to anti-drug antibody (ADA) formation including low detection rates in the clinic^6,10,14,18^, rapid onset compared to typical ADA timescales^20–22^, and a direct link between IL-12 receptor binding and desensitization^18^. For PK desensitization, the primary molecular mechanisms discussed revolve around the upregulation of IL-12 receptors on T-cells following IL-12 stimulation^14,23^. Increased expression of IL-12 receptor leads to higher levels of receptor-mediated clearance and faster serum clearance or increased IL-12 sequestration during transport into the blood. Overall, this would lead to reduced serum IL-12 concentrations due to increased clearance or lower bioavailability, and these effects are apparent almost immediately after repeated doses. Conversely, desensitization of IL-12’s immunological effects is delayed and tends to occur after several doses have been administered^14^. Discussed mechanisms for this effect include downregulation of STAT4 after sustained IL-12 signaling^24^ and adaptive T-cell exhaustion due to IL-12 stimulation^9,17^.

While no IL-12 therapies have been yet approved, many IL-12 technologies are in development with over 10 active or recruiting clinical trials on ClincialTrial.gov and several other technologies in preclinical development^25–33^. Due to its implications in both tolerability and efficacy of IL-12 treatment, understanding IL-12 desensitization is essential in the successful development of IL-12-based therapies. Here, we investigated mechanisms for IL-12 PK desensitization via a modeling approach. We developed mathematical models containing two different mechanisms of interest for IL-12 PK desensitization and challenged these models to fit clinical trial data. After identifying the mechanism that is consistent with the data, we determine the parameter regime under which significant desensitization can occur and use the model with our proposed mechanism to identify potential strategies for mitigating systemic toxicity to IL-12 treatment.

## Methods

### Clinical Trial Data Collection

To validate the models used in this investigation, we conducted a literature search to identify clinical trials satisfying two conditions: i) data (time-course or PK metrics) describing IL-12 PK over multiple doses are provided, and ii) clear evidence of IL-12 PK desensitization is present. Three clinical trials satisfying these conditions were identified (**Table S1**) ^14–16,18^. In addition to satisfying the conditions, these three clinical trials were determined to be well suited for this investigation because they provide IL-12 PK data after repeated doses of IL-12 with either different dosing regimens or doses selected for PK analysis (**Figure 1A**). Two of the clinical trials investigated report IL-12 time-course measurements, whereas the other trial only reports PK metrics, including area under the curve (AUC), maximum concentration (C_max_), and time of maximum concentration (t_max_). In trials with time-course data, the numerical values were extracted from figures using GraphGrabber2.0.2 (Quintessa).

**Figure 1:**
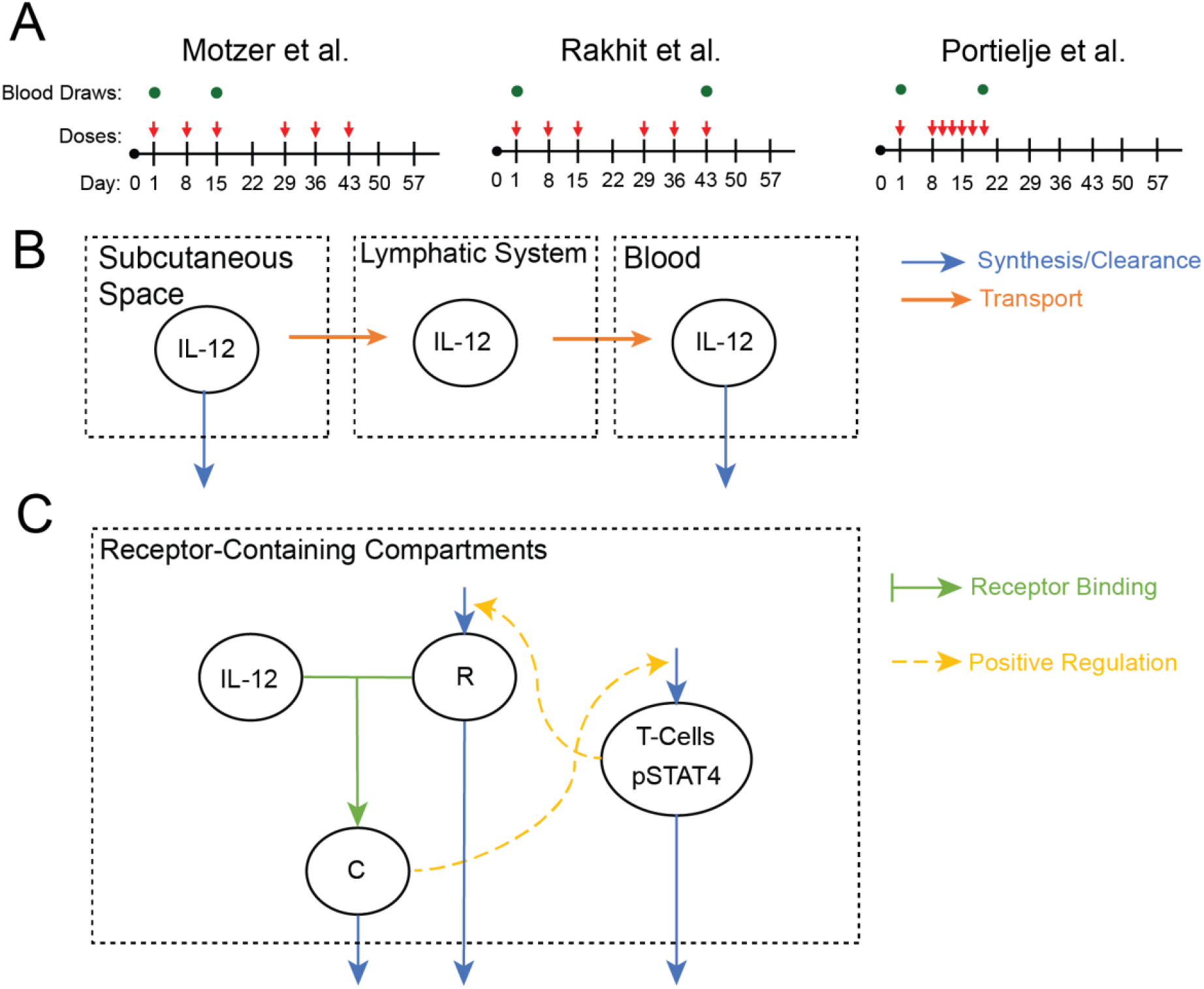
Clinical trial dosing information and schematics of models used in this investigation. A) Timelines illustrating dose regimens and doses where blood measurements are taken in three clinical trials used in this investigation. B) Schematic of IL-12 transport. C) Schematic of IL-12 receptor binding and upregulation.

### Mathematical Models of IL-12 PK and Numerical Simulation

The absorption, distribution, metabolism, and elimination (ADME) properties of IL-12 are not characterized in detail, but the large molecular weight of IL-12 (70 kDa) provides insight into its absorption and elimination characteristics. As molecules with greater molecular weights than 16 kDa are absorbed into the blood primarily by the lymphatic system^34^, it follows that IL-12 likely is absorbed via the same route following administration into peripheral tissues. Similarly, molecules with molecular weights over 50 kDa are typically eliminated via intracellular lysosomal proteolytic degradation rather than renal clearance^35,36^. However, renal clearance may still influence IL-12 elimination^37^.

Based on these ADME properties, we developed two mathematical models representing IL-12 dosing, transport, target binding and upregulation, and clearance. The notation and the meaning for each model parameter are found in **Table S2**. Below, we briefly explain the differential equations; derivations are provided in the **SI Text**.

Both models include identical equations governing IL-12 transport and non-receptor mediated clearance. IL-12 transport from the subcutaneous space, through the lymphatic system, and into the blood via lymph flow **(Figure 1B**) are described by:

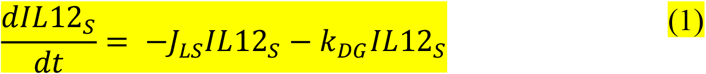

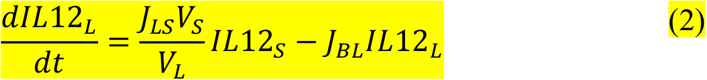

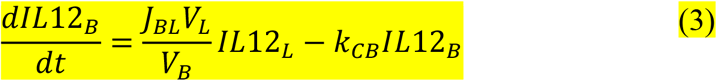

where *IL*12_*i*_ is the concentration of IL-12 in compartment *i* (pmol mL^-1^), *V*_*i*_is the volume of compartment *i* (mL), *J*_*ij*_ is the first-order transport rate constant from compartment *j* to compartment *i, k*_*DG*_is the first-order degradation rate constant for loss of IL-12 in the subcutaneous space (days^-1^), and *k*_*CB*_ is the first-order systemic clearance rate constant (days^-1^). Degradation in the subcutaneous space is related to the absolute bioavailability, *F*:

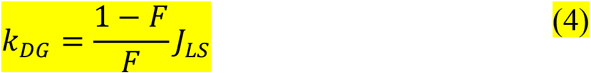

This relationship is derived such that 100% bioavailability leads to a degradation rate constant of zero and vice versa. Bolus IL-12 injection is represented by the initial concentration of IL-12 determined by dose amount (*D*) and dose compartment volume:

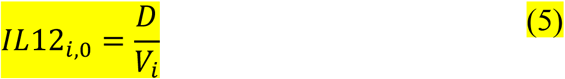

In addition to these terms governing transport of IL-12, both models developed contain IL-12 receptor binding and upregulation (**Figure 1C**). Receptor binding and STAT4 activation occurs in either the lymphatic system or the blood, and the concentration of IL-12:receptor complex is described as follows:

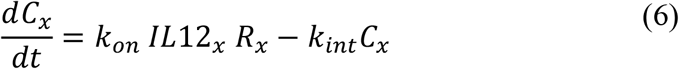

The first term represents the rate of IL-12:receptor binding, assumed to be irreversible, and the second term describes the first-order internalization of the complex. Formation of complex leads to STAT4 activation that is modeled by the following expression that assumes non-cooperative binding and a quasi-steady state:

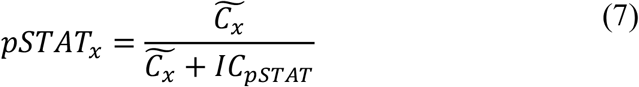

Here 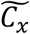 is the number of complexes per cell because STAT4 is activated on a single cell basis. 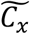 is calculated from the total concentration of complexes by assuming homogenous distribution:

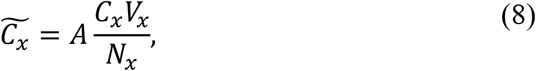

This equation converts the concentration of receptors in the compartment (pmol mL^-1^) to the number of complexes per T-cell (# cells^-1^) using Avogadro’s number 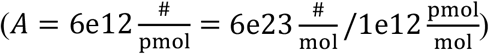, the volume of the compartment, and the number of T-cells in the compartment. In our models, activation of STAT4 leads to upregulation of IL-12 receptor on T-cells^15,18,23^. Accordingly, the concentration of IL-12 receptor in compartments is represented by:

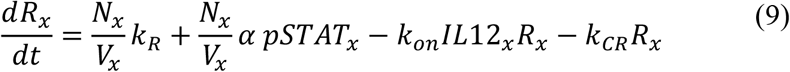

The first two terms in this ordinary differential equation represent basal and upregulated IL-12 receptor expression, respectively. The production rates, *k*_*R*_ and *α*, are given in units of pmol cells^-1^; thus, we scale by the number of cells divided by the volume to convert to pmol mL^-1^ days^-1^. The third term represents consumption of IL-12 receptor due to binding, and the last term represents unbound internalization of IL-12 receptor.

The initial concentration of receptor in compartment *x, R*_*SS*.*x*_ (pmol mL^-1^), is determined by *R*_0_, the basal amount of IL-12 receptor on the surface of T-cells (pmol), as follows:

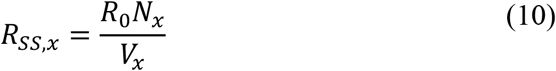

To ensure that this value is consistent with our model in the absence of IL-12, we constrain the basal production rate according to:

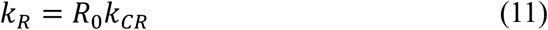

All models in this investigation were constructed using this framework, differing in the compartment in which receptor binding takes place. Models were simulated in MATLAB (R2020b) using the ODE solver ode15s. All option arguments are set to default except for the non-negative argument set to one and the absolute tolerance set to 1e-12.

### Data Fitting and Parameter Estimation Procedures

Model parameters were estimated by optimizing model fitness to data from each of the three clinical trials individually. Optimization boundaries are in **Table S3**. Models were fitted to PK datasets separately to determine the influence of dose regimen, sampling times, and data type on the variability of PK parameters. Two optimization methods were used: global optimization was implemented using MATLAB’s particleswarm algorithm, and ensemble local optimization was implemented by randomly sampling parameter sets within optimization boundaries and locally optimizing using MATLAB’s patternsearch algorithm^38^. The global optimization method provides a single, best-fitting parameter set whereas the ensemble local optimization method provides distributions of parameter values to account for variability in the datasets used. The fitness functions used for optimization were modified to account for assay quantification limits and varied based on the type of PK data presented; for specific details, see the supplementary text.

## Results

### Two alternative models of desensitization describing literature-proposed mechanisms were developed

We first identified proposed mechanisms for IL-12 PK desensitization in literature as a basis for developing our mathematical models. Analysis of preclinical and clinical trials of IL-12 provided evidence for two distinct causes of IL-12 PK desensitization: increased clearance or decreased bioavailability following repeated doses^10,14,15,18^. While both were previously proposed, neither cause has been validated. Thus, we aimed to formulate molecular mechanisms for the identified causes of IL-12 desensitization in literature and determine the likelihood for these mechanisms to account for IL-12 PK desensitization based on their ability to fit clinical trial data.

Two models were constructed based on the framework described in methods to describe molecular mechanisms for increased clearance or reduced bioavailability after repeated doses. Our first model, the accelerated-clearance model (**Equations S1-10**) only contains IL-12 receptor interaction in the blood such that IL-12 clearance from the blood is increased when receptor upregulation occurs. Conversely, the reduced-bioavailability model (**Equations S11-20**) only contains IL-12 receptor interaction in the lymphatic system. Here, IL-12 receptor upregulation leads to increased receptor interaction during transport to the blood, leading to sequestration. These two models are effectively limiting cases of the same model where receptor binding occurs in both the blood and lymphatic system with the number of receptor-expressing cells in one compartment set to zero. These limiting cases were analyzed to assess the likelihood of each mechanism independently to determine if either is capable of being solely responsible for IL-12 PK desensitization.

### The accelerated-clearance model fails to accurately predict IL-12 PK following repeated doses

To understand whether the accelerated-clearance model can capture trends of IL-12 desensitization in published data, we fitted the model to three separate clinical PK datasets of repeated subcutaneous IL-12 injections. The results are presented in **Figures 2A-C** (see also: **Tables S4-5**). The accelerated-clearance model can predict the presence of IL-12 PK desensitization; however, the level of receptor upregulation required for desensitization to occur leads to inaccurate clearance kinetics. This is especially clear when simulating repeated i.v. doses, where this model predicts a time dependent clearance rate that is inconsistent with clinical data **(Figure S1A**) ^6,7,13^. Therefore, we conclude that IL-12 receptor upregulation in the blood is unlikely to have a significant impact on IL-12 PK and lead to desensitization. Accordingly, the proposed accelerated-clearance model is unable to accurately predict IL-12 PK.

**Figure 2:**
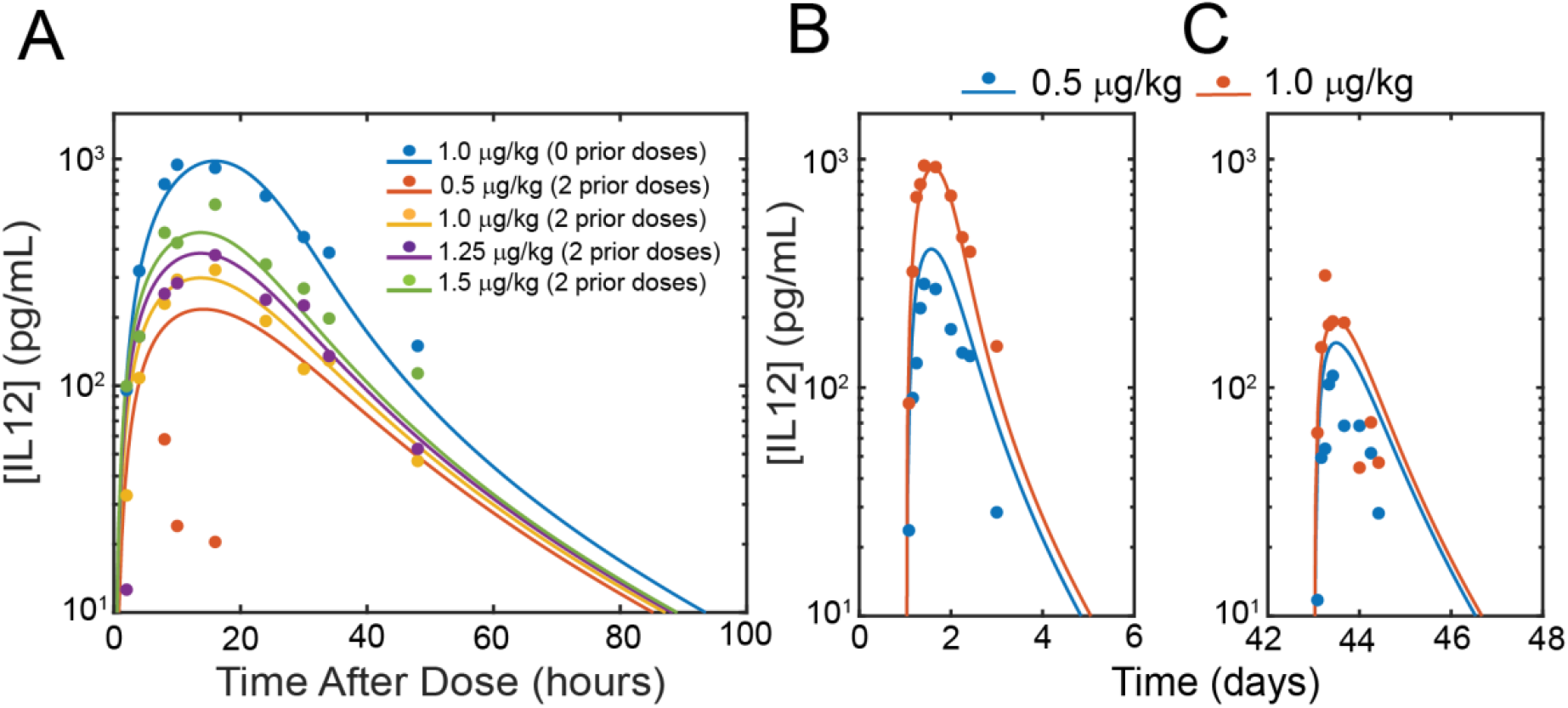
Accelerated-clearance model fitness to clinical trial data. A) Accelerated-clearance model fit to clinical trial data presented by Motzer et al. B-C) Accelerated-clearance model fit to clinical trial data presented by Rakhit et al. B) Predictions and data after first dose. C) Predictions and data after sixth dose.

We next aimed to further assess whether changes in clearance are consistent with the quantitative behavior of IL-12 PK desensitization. To this end, we removed feedback from the accelerated-clearance model (see equation S4), and separately fit IL-12 PK data presented by Motzer and colleagues after 1.0 µg/kg doses with and without prior exposure to IL-12. We first fit the dataset without prior exposure to estimate the model parameters (**Figure S2A**) and then fit the dataset with previous exposure by varying either the IL-12 clearance rate (*k*_*CB*_) or IL-12 bioavailability and absorption (*J*_*LS*_ and *F*) and keeping all other parameters fixed. As our fitting results suggested, the trends in data are inconsistent with changes in IL-12 clearance kinetics (**Figure S2B**). However, by manipulating bioavailability and absorption, the clearance kinetics can be accurately matched. This further supports the conclusion that increases in systemic clearance are not primarily responsible for IL-12 PK desensitization.

### The reduced-bioavailability model accurately predicts IL-12 PK across all three clinical trials

Next, we sought to determine whether our proposed mechanism influencing IL-12 bioavailability and transport into the blood can accurately fit IL-12 PK data reported in clinical trials. To this end, we fit the reduced-bioavailability model to all three clinical trials via global optimization. Optimization results illustrate that this model can fit these datasets accurately (**Figures 3A-C, Figure S3-4, Tables S6-7**). Moreover, unlike the accelerated-clearance model, the reduced-bioavailability model makes accurate predictions regarding i.v. dosing (**Figure S1B**). However, the 0.5 μg/kg dose in the dataset presented by Motzer et al.^14^ is not fit accurately. There is nonlinearity in the dataset that leads to concentrations at this dose level to be overpredicted in this parameter regime. Further analyses illustrate parameter sets with slower IL-12 transport from the lymphatic system to the blood (decreasing *J*_*BL*_) and stronger STAT4 activation (decreasing *IC*_*pSTAT*_) can capture this nonlinearity more accurately (**Figure S5, Table S8**), but fitness to high dose data is worsened. Given that the 0.5 µg/kg dataset presented is far below the maximum tolerated dose of 1.5 µg/kg and exhibited negligible toxicity and efficacy^14^, we decided to prioritize model parameter sets that more accurately fit the clinically relevant doses for the remainder of the paper.

**Figure 3:**
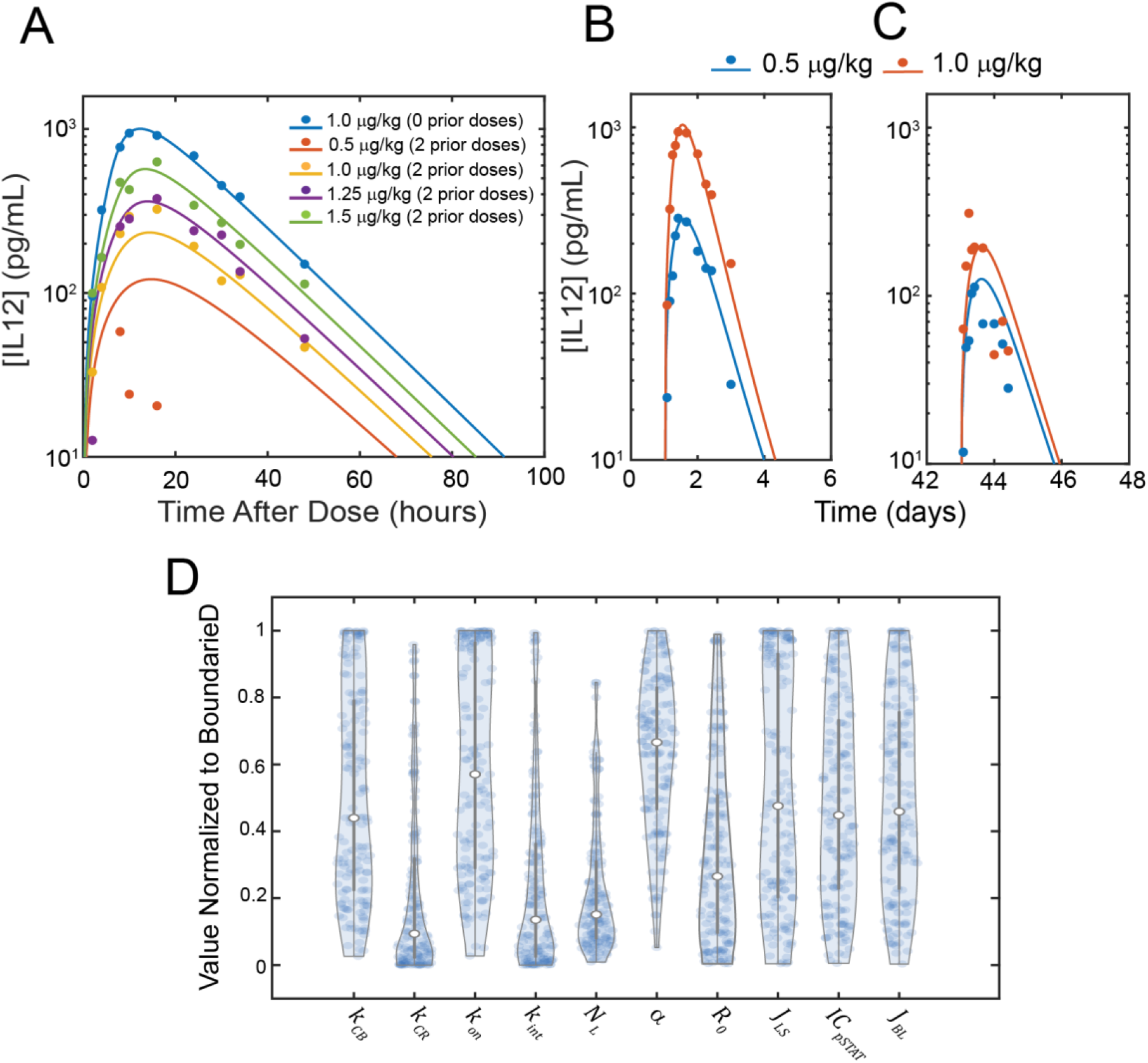
Reduced-bioavailability model fit to clinical trial data. A) Reduced-bioavailability model fit to clinical trial data presented by Motzer et al. after global optimization. B-C) Reduced-bioavailability model fit to clinical trial data presented by Rakhit et al. after global optimization B) Predictions and data after first dose. C) Predictions and data after sixth dose. D) Distribution of generated ensemble fit to all three clinical trials following local optimization of randomly sampled parameter sets within boundaries (50 parameter sets per clinical trial).

As PK parameters often vary between patients^39^, we sought to determine if our model can accurately fit IL-12 PK data over a wide range of biologically relevant parameter values. An ensemble of parameter sets was obtained by locally optimizing many sampled parameter sets to each clinical trial. This method yields parameter sets with slightly worse fitness to data when compared to global optimization (**Figure S6, Table S9**) but gives us a distribution of parameter sets to analyze. Analysis of parameter distributions from all three fits illustrates that model parameters are widely distributed over the optimization boundaries (**Figure 3D**). Furthermore, the distribution of parameter values does not vary significantly based on the clinical trial used for fitting (**Figure S7**). These results illustrate that the ability of our model to predict IL-12 PK desensitization is not restricted to small regions of the parameter space, suggesting that IL-12 PK desensitization is not overly sensitive to interpatient variations in PK parameters. Additionally, the PK datasets available are insufficient to fully constrain PK parameters.

### Analysis of model behavior illustrates the effects of dosing regimen and model parameters on desensitization

To gain insight into how the dosing regimen and model parameters influence the amount of IL-12 PK desensitization in the reduced-bioavailability model, we simulated the model under various dosing conditions and parameter values to determine the effects on PK desensitization. For this analysis we use the parameters obtained via global optimization to the data presented by Motzer and colleagues^14^. This parameter set was selected because fitness resulted in a lower squared error compared to the fit to the data presented by Rakhit et al.^18^ or both simultaneously (1.80e-5 over 41 datapoints v. 2.20e-5 over 38 datapoints v. 3.86e-5 over 79 datapoints) and the dataset used for fitting is more detailed than the Portielje et al. dataset^15^. We first investigated the amount of desensitization our model predicts following two consecutive doses of IL-12 with varying dose amounts (**Figure 4A**) and time between doses (**Figure 4B**). Predictions illustrate that desensitization increases with higher doses and doses that are closer in time. We hypothesized that desensitization is mainly controlled by the concentration of receptor remaining in the lymphatic system by the time of the next dose. A larger dose leads to increased STAT4 activation and a more significant increase in receptor expression, and doses closer in proximity lead to less receptor clearance back to baseline before the next dose. To expand on this hypothesis, we generated an ensemble of 10,000 parameter sets varying *k*_*CR*_, *k*_*int*_, *N*_*cells*_, *α, R*_0_, and *IC*_*pSTAT*_ between 50% and 150% of the optimized values holding all other parameters constant and simulated consecutive s.c. doses of IL-12 one week apart. Analysis of the ensemble simulation results validated our hypothesis that the extent of desensitization predicted correlates with the difference in receptor concentrations between the first and second dose and the sensitivity of IL-12 PK to receptor binding (**Figure S8**).

**Figure 4:**
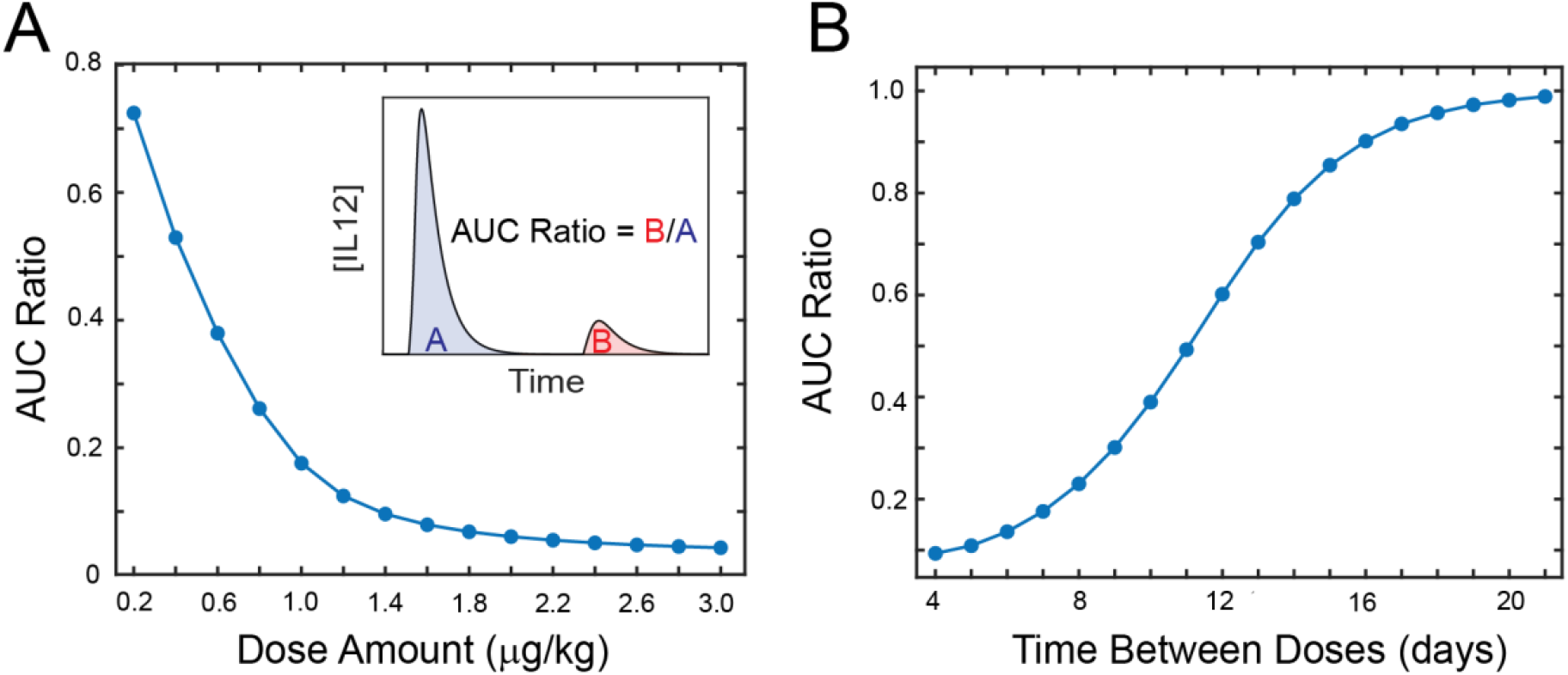
Analysis of dosing regimen on predicted amounts of desensitization. A) Predicted amount of desensitization after two consecutive IL-12 doses seven days apart at varying dose levels. Inset: visual representation of AUC ratio. B) Predicted amount of desensitization after two consecutive 1.0 µg/kg doses with varying time between doses.

We next analyzed how variations in specific parameters related to receptor upregulation influence PK desensitization. Local sensitivity analyses were implemented for two and three consecutive doses of varying amounts (**Figure S9**). The only model parameter with negligible impact on desensitization was *k*_*CB*_. Of interest, parameters related to the amount of receptor initially (*R*_0_, *N*_*cells*_) and the lifetime of receptor (*k*_*CR*_, *k*_*int*_) had log gain sensitivities greater than one at some dose levels.

### Simulation of local IL-12 delivery reveals potential strategies for limiting systemic exposure to IL-12 while maintaining desired local concentrations

Next, we sought to explore the consequences of our proposed mechanism in designing IL-12 therapies for limiting systemic exposure to IL-12. As current interest in IL-12 therapy revolves around local cytokine delivery via cell or gene therapy^8,9,12,17,40,41^, we modified the reduced-bioavailability model to represent bolus or continuous delivery of IL-12 to the peritoneal cavity (**Figure S10, Equations S21-S39**). This model is structurally identical to the reduced-bioavailability model, except the volume of the dosing compartment is increased to the volume of the peritoneal cavity, the transport parameters are re-estimated, and a 0^th^ order intraperitoneal IL-12 production term is introduced for simulation of continuous delivery. We selected intraperitoneal administration for these analyses because it is a common site of administration for IL-12-based therapies^42–44^, and a PK dataset is available reporting both intraperitoneal and serum IL-12 concentrations following intraperitoneal IL-12 injection. Using this dataset to estimate transport-related parameters allows us to have greater confidence in both our predicted serum and intraperitoneal IL-12 concentrations. Accordingly, we first estimated the parameters describing IL-12 transport from the intraperitoneal space to the blood by fitting our model to clinical trial data presented by Lenzi et al.^45^ modified to represent intraperitoneal injection of IL-12. Our model was able to accurately fit both the intraperitoneal and serum IL-12 concentrations (**Figure S11, Table S10**), giving us confidence in the estimated transport parameters. Additionally, a standard PK model is unable to fit this data, further supporting the relevance of our model. Since this dataset only consisted of a single intraperitoneal infusion of IL-12, we used the transport parameters estimated from the Lenzi et al. data^45^ and parameters to characterize receptor-upregulation feedback estimated from our fits in earlier sections (**Table S11**).

We next used this model to simulate continuous IL-12 delivery to the peritoneal cavity and predicted intraperitoneal and serum concentrations of IL-12 according to three different production profiles: 1) constant production, 2) step-wise production, and 3) linearly-increasing production designed to reach an equivalent concentration of IL-12 in the intraperitoneal space at steady state (**Figure 5A**). Importantly, our model predicts serum IL-12 concentrations exceeding levels where toxicity is observed^14^ following the constant production profile, whereas both of the dynamic dosing strategies maintain serum concentrations below the healthy maximum^46^. These results suggest that systemic exposure to IL-12 may be avoided following local delivery by utilizing dosing strategies that slowly reach the desired production rate for local concentrations rather than immediately starting at that level.

**Figure 5:**
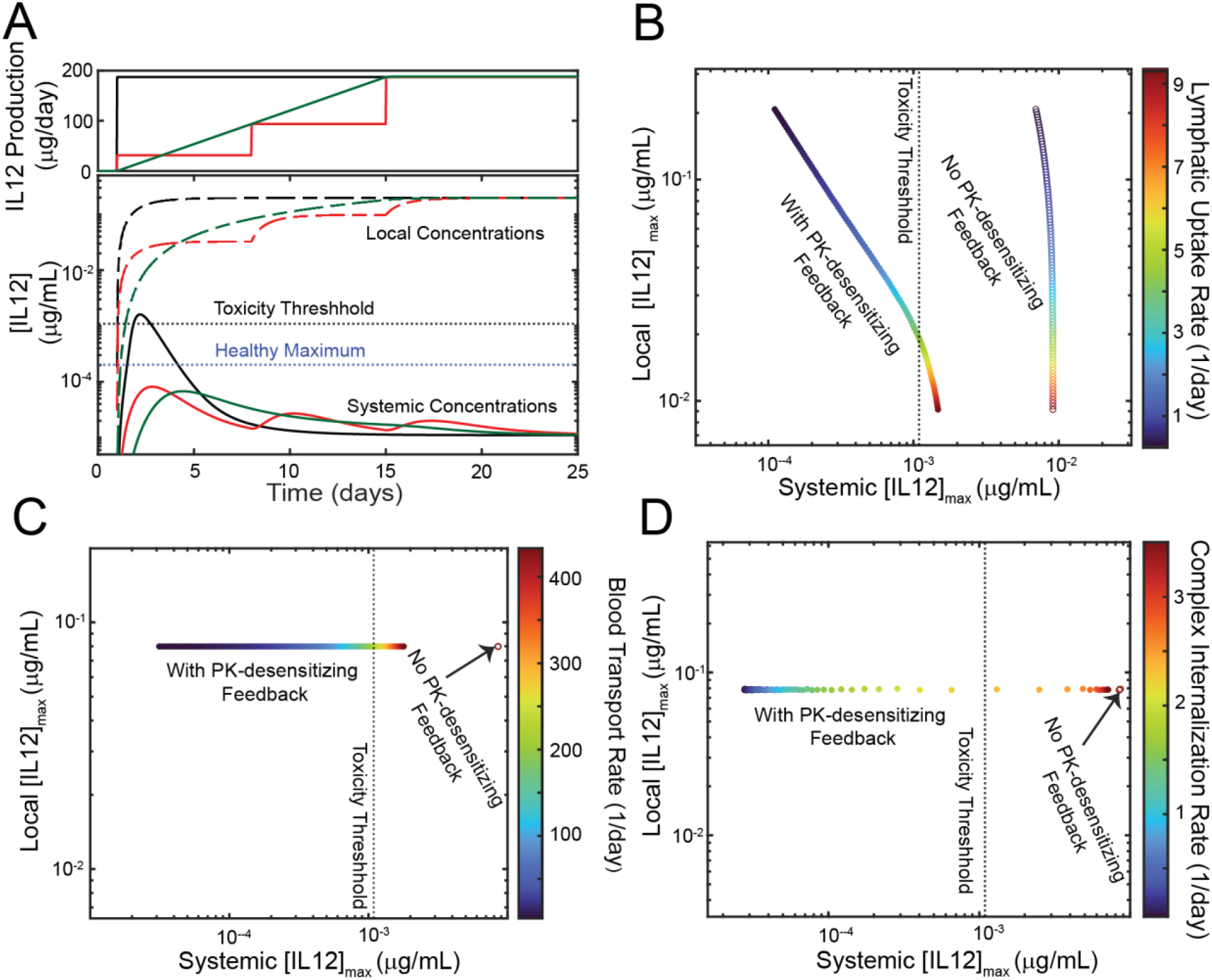
Model predictions of strategies for mitigating systemic IL-12 exposure following continuous local delivery. A) Prediction of continuous, intraperitoneal IL-12 delivery according to three different production rate profiles: constant production (black lines), step-wise production (red lines), and linearly increasing production (green lines). Healthy IL-12 serum concentration taken from Derin et al.^46^ B-D) Effects of various IL-12 related parameters on maximum concentrations in the local environment and in the blood. B) Effects of manipulating lymphatic uptake. C) Effects of limiting transport into the blood from the lymphatic system. C) Effects of limiting internalization of IL-12:IL-12 receptor complex.

In addition to analyzing the predicted effects of dynamic dosing strategies, we used our model to identify potential protein engineering approaches for limiting systemic exposure. Based on our simulations, strategies modifying IL-12 to limit uptake into the lymphatic system from the local environment (**Figure 5B**), limit uptake into the blood from the lymphatic system (**Figure 5C**), or limit internalization of IL-12:receptor complex (**Figure 5D**) were predicted to have profound effects due to our mechanism. We found that these aspects of IL-12 could be manipulated such that the systemic exposure to IL-12 is reduced, and the concentrations in the local environment are either increased or unchanged. These strategies are a direct result of IL-12 PK desensitization; without our mechanism, they are not predicted to be effective. Ultimately, these results illustrate how our proposed mechanism for IL-12 desensitization can be leveraged to identify strategies for limiting systemic exposure to IL-12 treatment.

## Discussion

Here, we employ a modeling approach to assess the relevance of proposed mechanisms for IL-12 PK desensitization. Two models of IL-12 PK containing feedback mechanisms causing increased clearance or reduced bioavailability following repeated doses of IL-12 were developed. We found that the level of receptor upregulation/increase in IL-12 clearance required for the accelerated-clearance model to predict desensitization leads to inaccurate clearance kinetics and inconsistent predictions of i.v. doses. Conversely, the reduced-bioavailability model accurately fit IL-12 PK across three separate clinical trials and correctly predicts little desensitization with repeated i.v. doses^6,7,13^. These results provide quantitative evidence that IL-12 PK desensitization results from decreased bioavailability with repeated doses rather than increased clearance. Additionally, we developed a model framework that can accurately predict IL-12 PK over repeated doses, which is not currently available to the best of our knowledge.

The proposed molecular mechanism for IL-12 PK desensitization is that IL-12 stimulation of T-cells in the lymphatic system leads to increased expression of the IL-12 receptor and thereby increased sequestration during transport into the blood. Upregulation of IL-12 receptor following IL-12 activity is shown both *in vitro* and *in vivo* in the literature^18,23^. Our results provide quantitative justification for how upregulation of IL-12 receptor affects IL-12 PK. These results remain a hypothetical but experimentally testable prediction; however, we believe there is evidence that supports that this proposed mechanism for IL-12 PK is biologically feasible. Because IL-12’s molecular weight is greater than 16 kDa, it cannot enter the blood through capillary diffusion and must travel through the lymphatic system to enter the blood from peripheral tissue^34^. T-cells are highly abundant in the lymphatic system^47^; therefore, during lymphatic transport, it is expected that significant receptor interaction will occur. Furthermore, our hypothesis requires interaction between IL-12 and its receptor for desensitization to occur. Rakhit and colleagues illustrate that knocking out the IL-12 receptor in mice prevents IL-12 desensitization^18^, illustrating that PK desensitization does not occur in the absence of receptor binding. Our mechanism also provides a plausible explanation for why PK desensitization may not occur in some cases. For example, studies of repeated i.v. administration demonstrate that negligible desensitization occurs^7,13^, suggesting that the responsible mechanism is more significant in peripheral tissues. Additionally, Ohno et al.^48^ report IL-12 PK following repeated subcutaneous doses with a maximum dose of 3 µg/kg where PK desensitization is not clearly observed. We demonstrated that our hypothesis predicts that IL-12 PK desensitization is dose-dependent, which could perhaps have a threshold level of activation for occurring that is not included in our model. Altogether, we believe that our understanding of large molecule PK and evidence in literature suggests our proposed mechanism is highly realistic.

Due to limitations in the datasets available for model validation, model parameters pertaining to molecular details of the proposed feedback mechanism could not be estimated with high confidence. We instead demonstrated that while specific parameter values were not tightly constrained, the ability of our model to predict PK desensitization was related to the amount of receptor upregulation between doses and model sensitivity to lymphatic receptor concentrations. While this does not allow us to specifically determine model parameters, the primary outcome of this investigation is validation that upregulation of IL-12 receptor during lymphatic transport is a feasible explanation for IL-12 PK desensitization. This conclusion is not dependent on parameter identifiability. Furthermore, the ability of a wide range of model parameters to predict desensitization suggests that this mechanism can be relevant in the context of other biologics as well. Any biologic that i) upregulates the expression of its receptor, ii) enters the blood via lymphatic transport following extravascular dosing, and iii) binds a target with high abundance in the lymphatic system may exhibit similar effects. A potential example is pegfilgrastim, an FDA-approved granulocyte colony-stimulating factor^49,50^.

Despite decades of failure in the clinic, interest in developing IL-12-based therapies remains high with many technologies in clinical or preclinical development^25–33^. The development of IL-12 therapies today focuses on local delivery of IL-12 through gene or cell-based therapies to minimize systemic exposure to IL-12^8,9,12,17,40,41^. Importantly, technologies where significant IL-12 exposure occurs cause severe toxicities, limiting the therapeutic window of treatment^41^. Due to the inherent role of IL-12 PK desensitization on systemic exposure to IL-12, our results will undoubtedly aid in the modern development of IL-12-based therapies. The final set of analyses presented in this investigation quantitatively illustrate how IL-12 PK desensitization can be leveraged to maintain high local concentrations of IL-12 while minimizing systemic exposure. The effects of variable IL-12 production rates and protein engineering strategies are illustrated, which are not obvious to be effective without use of our model. Beyond the analyses completed in this investigation, we provide a mathematical modeling framework that can be applied to future development of IL-12-based therapies.

## Study Highlights

### What is the current knowledge on this topic?

IL-12 desensitization was observed in the earliest clinical trials investigating IL-12 immunotherapy, and several studies have linked the phenomenon to treatment efficacy and tolerability. Mechanisms have been hypothesized in literature, but none have been validated quantitatively.

### What question did this study address?

What is the desensitization mechanism capable of correctly predicting IL-12 pharmacokinetics over repeated doses, and what are the therapeutic consequences of that mechanism?

### What does this study add to our knowledge?

Here, we developed a mathematical model that attempts to validate the mechanisms causing IL-12 PK desensitization in the absence of mechanistic data. Our results illustrate that IL-12 PK desensitization is likely to be a result of decreased bioavailability after repeated exposure to IL-12 rather than increased systemic clearance. We propose a molecular mechanism for this model based on the upregulation of IL-12 receptor on T-cells in the lymphatic system.

### How might this change drug discovery, development, and/or therapeutics?

Pharmacokinetic modeling is essential in predicting the behavior of therapeutics in humans. We provide a tool for predicting IL-12 pharmacokinetics, which will undoubtedly aid in the development of IL-12 based therapies.

## Author Contributions

J.D., O.A.I., and O.V. wrote the manuscript; J.D. and O.A.I. designed the research; J.D performed the research; J.D. and O.A.I. analyzed the data.

## Supplementary Material

**Supplementary Table S1:** Information on clinical trials analyzed in this investigation.

| **Clinical Trial** | **Dose Amounts and Regimen** | **Number of Patients** | **Patient Indications** | **Sampling Times** |
| --- | --- | --- | --- | --- |
| Phase I trial of Subcutaneous Recombinant Human Interleukin-12 in Patients with Advanced Renal Cell Carcinoma<sup>1</sup> | 0.1 – 1.5 µg/kg on days 1, 8, and 15 in two different 28-day cycles. Cycle 1: Fixed cycle with constant dose given all three days. Cycle 2: Escalating cycle with incrementally increasing doses until target dose is reached on day 15. | 51 total patients. Cycle 1, 0.5 µg/kg: 15 patients; Cycle 1, 1.0 µg/kg: 6 patients; Cycle 2, 0.5 µg/kg target dose: 3 patients; Cycle 2, 1.0 µg/kg target dose: 3 patients; Cycle 2, 1.25 µg/kg target dose: 6 patients; Cycle 2, 1.5 µg/kg target dose: 12 patients. | Advanced renal cell carcinoma. 42 patients received prior therapy. | Baseline and 10, 24, 48, 72, 96, and 168 h after dosing. PK time course data reported for dose 1 in cycle 1 and dose 3 in cycle 2. |
| Phase I Study of Subcutaneously Administered Recombinant Human Interleukin 12 in Patients with Advanced Renal Cell Cancer<sup>2</sup> | 0.1, 0.5, 1.0, 1.25 µg/kg given as a single dose or three times weekly for two weeks following a test dose one week prior for studying repeat administration. | 28 total patients initially enrolled, 25 patients included in data analysis. PK dataset includes single dose data for 0.5 µg/kg (n = 12), for 1.0 µg/kg (n = 4) and repeat dose data for 0.5 µg/kg (n = 4) and 1.0 µg/kg (n = 2). | Advanced renal cell carcinoma. 26 patients received prior therapy. | Baseline and 4, 6, 8, 10, 12, 18, 24, 30, 26, 48, 72, 96 h after test dose and 4, 6, 8, 10, 12, 18, 24, and 36 h after repeat doses. PK metrics are reported for doses one and seven following 0.5 and 1.0 µg/kg doses. |
| Down-regulation of the pharmacokinetic-pharmacodynamic response to interleukin-12 during long-term administration to patients with renal cell carcinoma and evaluation of the mechanism of this “adaptive response” in mice<sup>3</sup> | 0.1, 0.5, 1.0 µg/kg on days 1, 8, and 15 of a 28-day cycle. | 24 total patients. 3 patients given 0.1 µg/kg doses, 15 given 0.5 µg/kg doses, and 6 patients given 1.0 µg/kg doses. | Advanced or metastatic renal cell carcinoma. 15 patients received prior therapy. | Baseline and 2, 4, 8, 8, 10, 16, 24, 30, 34, 48 h post dose on dose 1 of cycle 1 and dose 3 of cycle 2. PK time course data reported for dose one of first cycle and dose three of second cycle for all dose levels. Reported concentrations for 0.1 µg/kg are below assay limit of detection so omitted from analysis. |

**Supplementary Table S2:** General model parameters, species, and meanings.

| Species | Meaning (Units) |
| --- | --- |
| $IL12_x$ | Concentration of IL-12 in compartment $x$ (pmol mL<sup>-1</sup>). |
| $R_x$ | Concentration of IL-12 receptor in compartment $x$ (pmol mL<sup>-1</sup>) |
| $C_x$ | Concentration of IL-12:IL-12 receptor in compartment $x$ (pmol mL<sup>-1</sup>). |
| $\widetilde{C}_x$ | Number of IL-12:IL-12 receptor complexes per T-cell in compartment $x$ (# cells<sup>-1</sup>) |
| $pSTAT_x$ | Fractional of maximum STAT4 activation on T-cells (unitless) |
| Parameter | Meaning (Units) |
| $D$ | IL-12 dose administered (pmol). |
| $J_{xy}$ | Normalized lymphatic fluid flow rate from compartment $y$ to compartment $x$ (days<sup>-1</sup>). |
| $k_{DG}$ | First order IL-12 degradation rate constant in the dosing compartment (days<sup>-1</sup>) |
| $F$ | Absolute IL-12 bioavailability defined as fraction of IL-12 leaving the dosing compartment instead of being degraded (unitless) |
| $k_{CB}$ | First order IL-12 systemic clearance rate constant from blood compartment (days<sup>-1</sup>) |
| $k_{on}$ | IL-12:IL-12 receptor association rate constant (mL pmol<sup>-1</sup> days<sup>-1</sup>) |
| $k_{int}$ | IL-12 receptor bound internalization rate constant (days<sup>-1</sup>) |
| $k_{CR}$ | IL-12 receptor unbound internalization rate constant (days<sup>-1</sup>) |
| $IC_{pSTAT}$ | Number of IL-12:IL-12 receptor complexes per T-cell required for half-maximum STAT4 activation (# cells<sup>-1</sup>) |
| $k_R$ | Basal IL-12 receptor production rate per T-cell (pmol cells<sup>-1</sup> days<sup>-1</sup>) |
| $\alpha$ | Receptor upregulation feedback coefficient (pmol cells<sup>-1</sup> days<sup>-1</sup>) |
| $N_x$ | Number of T-cells in compartment $x$ (#) |
| $R_0$ | Basal amount of IL-12 receptor on T-cell (pmol cells<sup>-1</sup>) |
| $V_x$ | Volume of compartment $x$ (mL) |

**Supplementary Table S3:** Model parameter information and sources. AC = accelerated-clearance, RB = reduced-bioavailability.

| Parameter | Meaning | Optimization Range/Default Value | Sources |
| --- | --- | --- | --- |
| $J_{LS}(\text{days}^{-1})$ | First order IL-12 transport rate from subcutaneous space to lymphatic system. | 0.79 – 2.4 | Milewski et al.<sup>6</sup> |
| $J_{BL}(\text{days}^{-1})$ | First order IL-12 transport rate from lymphatic system to blood. | 4.3 – 430 | Bounds centered around value reported by Varkhede et al.<sup>7</sup> |
| $k_{CB}(\text{days}^{-1})$ | First order IL-12 clearance rate from blood. | 0.79 – 2.4 | Reported half-life values in clinical trials (Motzer et al.<sup>1</sup>, Rakhit et al.<sup>3</sup>, Portielje et al.<sup>2</sup>) |
| $k_{DG}(\text{days}^{-1})$ | First order degradation rate of IL-12 in the subcutaneous space. | n/a | Determined by equation S18. |
| $F(\text{unitless})$ | Absolute bioavailability, fraction of IL-12 reaching lymphatic system. | 0.01 – 1 (AC model)<br>1 (RB model) | Assumed to be one to isolate bioavailability mechanism. |
| $k_{on}(\text{mL pmol}^{-1}\text{days}^{-1})$ | IL-12:IL-12 receptor association rate. | 1.5 – 210 | Chizzonite et al.<sup>4</sup>, adjusted to have high binding rate for improved fit to data. |
| $k_R(\text{pmol cells}^{-1}\text{days}^{-1})$ | Basal receptor production rate. | n/a | Determined by equation S17. |
| $k_{CR}(\text{days}^{-1})$ | IL-12 receptor unbound internalization rate. | 0.46 – 1.7 | Literature estimate unavailable, estimated based on half life between 10 and 36 hours. |
| $k_{int}(\text{days}^{-1})$ | IL-12 receptor bound internalization rate. | 0.46 – 1.7 | Literature estimate unavailable, estimated based on half life between 10 and 36 hours. |
| $\alpha(\text{pmol cells}^{-1}\text{days}^{-1})$ | Linear feedback coefficient. | 1.0e-8 – 1.0e-5 | Estimated, given large range. |
| $IC_{pSTAT}(\# \text{ cells}^{-1})$ | Complex formation required for 50% maximum STAT4 activation. | $9.1\text{e}2 - 1.8\text{e}6$ | Estimated, given large range. |
| $N_L(\text{cells})$ | Number of IL-12R+ T-cells in the lymphatic system | $1.0\text{e}9 - 5.0\text{e}11$ | Sender et al.<sup>5</sup> |
| $N_B(\text{cells})$ | Number of IL-12R+ T-cells in the blood | $8\text{e}8 - 8\text{e}10$ | Sender et al.<sup>5</sup> |
| $IL12_{s,0}(\text{pmol mL}^{-1})$ | Initial concentration of IL-12 in the subcutaneous space. | n/a | Determined by equation S19. |
| $D(\text{pmol})$ | Dose of IL-12 administered | Variable based on clinical trial. | Based on clinical trial information. |
| $V_S(\text{mL})$ | Volume of injection in subcutaneous space. | Variable based on clinical trial. | Based on volume of injection for clinical trials. Assumed to be 1 mL if not specified |
| $V_L(\text{mL})$ | Volume of lymphatic system IL-12 enters in transport to blood. | 33 | Varkhede et al.<sup>7</sup> |
| $V_B(\text{mL})$ | Volume of blood. | 5100 | Average human blood volume. |

**Supplementary Table S4:** Reported IL-12 C_max_, AUC, and t_max_ by Portielje et al. following repeated s.c. injections compared to accelerated-clearance model predictions after global optimization. Predicted values are reported to two significant digits.

| Dose Information | Predicted $C_{max}$ | Reported $C_{max}$ | Predicted AUC | Reported AUC | Predicted $t_{max}$ | Reported $t_{max}$ |
| --- | --- | --- | --- | --- | --- | --- |
| 0.5 $\mu\text{g/kg}$, dose 1 | 430 pg/mL | 362 (214) pg/mL | 7,600 pg*h/mL | 7,400 (5,400) pg*h/mL | 9.1 h | 9.7 (5) h |
| 0.5 $\mu\text{g/kg}$, dose 7 | 240 pg/mL | 255 (200) pg/mL | 4,000 pg*h/mL | 3,300 (1,600) pg*h/mL | 7.4 h | 9.5 (2.2) h |
| 1.0 $\mu\text{g/kg}$,<br>dose 1 | 1,100<br>pg/mL | 1131<br>(1051)<br>pg/mL | 21,000<br>pg*h/mL | 31,800<br>(22,300)<br>pg*h/mL | 12 h | 17 (7) h |
| 1.0 $\mu\text{g/kg}$,<br>dose 7 | 350<br>pg/mL | 376 (49)<br>pg/mL | 5,900<br>pg*h/mL | 6,000<br>(2,400)<br>pg*h/mL | 7.6 h | 4 (0) h |

\*Values in () are reported standard deviation.

**Supplementary Table S5:** Accelerated-clearance model parameter values following global optimization to each clinical trial individually. Values are reported to two significant digits.

| Parameter | Value fit to<br>Motzer data | Value fit to Rakhit<br>data | Value fit to<br>Portielje data |
| --- | --- | --- | --- |
| $k_{CB}(\text{days}^{-1})$ | 2.3 | 2.3 | 0.8 |
| $k_{CR}(\text{days}^{-1})$ | 0.46 | 0.46 | 0.46 |
| $k_{on}(\text{mL pmol}^{-1}\text{days}^{-1})$ | 210 | 200 | 210 |
| $k_{int}(\text{days}^{-1})$ | 0.46 | 0.49 | 0.46 |
| $N_L(\text{cells})$ | 8.0e10 | 2.6e10 | 6.3e10 |
| $\alpha(\text{pmol cells}^{-1}\text{days}^{-1})$ | 08.8e-6 | 4.2e-6 | 2.6e-6 |
| $R_0(\text{pmol cells}^{-1})$ | 2.1e-9 | 8.1e-9 | 8.1e-9 |
| $J_{LS}(\text{days}^{-1})$ | 0.79 | 0.78 | 2.9 |
| $J_{BL}(\text{days}^{-1})$ | 4.33 | 6.2 | 4.4 |
| $F$ (unitless) | 1 | 1 | 0.77 |
| $IC_{pstat}(\# \text{ cells}^{-1})$ | 1.8e6 | 1.1e6 | 1.8e6 |

**Supplementary Table S6:** Reported IL-12 C_max_, AUC, and t_max_ by Portielje et al. following repeated s.c. injections compared to reduced-bioavailability model predictions after global optimization. Predicted values are reported to two significant digits.

| Dose<br>Information | Predicted<br>$C_{max}$ | Reported<br>$C_{max}$ | Predicted<br>AUC | Reported<br>AUC | Predicted<br>$t_{max}$ | Reported<br>$t_{max}$ |
| --- | --- | --- | --- | --- | --- | --- |
| 0.5 $\mu\text{g/kg}$,<br>dose 1 | 330<br>pg/mL | 362 (214)<br>pg/mL | 7,400<br>pg*h/mL | 7,400<br>(5,400)<br>pg*h/mL | 10 h | 9.7 (5) h |
| 0.5 $\mu\text{g/kg}$,<br>dose 7 | 170<br>pg/mL | 255 (200)<br>pg/mL | 4,100<br>pg*h/mL | 3,300<br>(1,600)<br>pg*h/mL | 10 h | 9.5 (2.2) h |
| 1.0 µg/kg,<br>dose 1 | 1,300<br>pg/mL | 1131<br>(1051)<br>pg/mL | 26,000<br>pg*h/mL | 31,800<br>(22,300)<br>pg*h/mL | 9.8 h | 17 (7) h |
| 1.0 µg/kg,<br>dose 7 | 290<br>pg/mL | 376 (49)<br>pg/mL | 6,800<br>pg*h/mL | 6,000<br>(2.4)<br>pg*h/mL | 10 h | 4 (0) h |

\*Values in () are reported standard deviation.

**Supplementary Table S7:** Reduced-bioavailability model parameter values following global optimization to each clinical trial individually and Motzer and Rakhit data simultaneously. Values are reported to two significant digits.

| Parameter | Value fit to<br>Motzer data | Value fit to<br>Rakhit data | Value fit to Motzer<br>and Rakhit Data | Value fit to<br>Portielje data |
| --- | --- | --- | --- | --- |
| $k_{CB}(\text{days}^{-1})$ | 1.6 | 2.4 | 2.0 | 2.4 |
| $k_{CR}(\text{days}^{-1})$ | 0.46 | 0.50 | 0.50 | 1.7 |
| $k_{on}(\text{mL pmol}^{-1}\text{days}^{-1})$ | 200 | 200 | 140 | 200 |
| $k_{int}(\text{days}^{-1})$ | 0.56 | 0.59 | 0.61 | 0.66 |
| $N_L(\text{cells})$ | 3.8e10 | 3.3e10 | 9.3e10 | 1.7e11 |
| $\alpha(\text{pmol cells}^{-1}\text{days}^{-1})$ | 5.0e-6 | 3.1e-6 | 6.7e-6 | 4.4e-6 |
| $R_0(\text{pmol cells}^{-1})$ | 5.6e-9 | 5.5e-9 | 1.7e-9 | 1.8e-9 |
| $J_{LS}(\text{days}^{-1})$ | 1.6 | 1.2 | 1.1 | 2.9 |
| $J_{BL}(\text{days}^{-1})$ | 70 | 130 | 70 | 150 |
| $IC_{pstat}(\# \text{ cells}^{-1})$ | 1.4e6 | 9.8e5 | 1.8e6 | 1.8e6 |

**Supplementary Table S8:** Reduced-bioavailability parameter set that predicts nonlinearity observed in Motzer data at expense of fitness to high dose dataset.

| Parameter | Value |
| --- | --- |
| $k_{CB}(\text{days}^{-1})$ | 1.3 |
| $k_{CR}(\text{days}^{-1})$ | 1.7 |
| $k_{on}(\text{mL pmol}^{-1}\text{days}^{-1})$ | 170 |
| $k_{int}(\text{days}^{-1})$ | 0.61 |
| $N_L(\text{cells})$ | 2.9e9 |
| $\alpha(\text{pmol cells}^{-1}\text{days}^{-1})$ | 5.7e-6 |
| $R_0(\text{pmol cells}^{-1})$ | 1.5e-8 |
| $J_{LS}(\text{days}^{-1})$ | 0.80 |
| $J_{BL}(\text{days}^{-1})$ | 12 |
| $IC_{pstat}(\# \text{ cells}^{-1})$ | 9.5e5 |

**Supplementary Table S9:** Reported IL-12 C_max_, AUC, and t_max_ by Portielje et al. following repeated s.c. injections compared to reduced-bioavailability model predictions after ensemble local optimization. Predicted values are reported to two significant digits.

| Dose Information | Predicted $C_{max}$ | Reported $C_{max}$ | Predicted AUC | Reported AUC | Predicted $t_{max}$ | Reported $t_{max}$ |
| --- | --- | --- | --- | --- | --- | --- |
| 0.5 $\mu\text{g/kg}$, dose 1 | 350 (30) pg/mL | 362 (214) pg/mL | 8,700 (570) pg*h/mL | 7,400 (5,400) pg*h/mL | 9.6 (1.1) h | 9.7 (5) h |
| 0.5 $\mu\text{g/kg}$, dose 7 | 170 (15) pg/mL | 255 (200) pg/mL | 4,500 (180) pg*h/mL | 3,300 (1,600) pg*h/mL | 9.7 (0.86) h | 9.5 (2.2) h |
| 1.0 $\mu\text{g/kg}$, dose 1 | 940 (190) pg/mL | 1131 (1051) pg/mL | 21,000 (4,200) pg*h/mL | 31,800 (22,300) pg*h/mL | 9.3 (1.1) h | 17 (7) h |
| 1.0 $\mu\text{g/kg}$, dose 7 | 240 (20) pg/mL | 376 (49) pg/mL | 6,300 (290) pg*h/mL | 6,000 (2,400) pg*h/mL | 9.8 (0.94) h | 4 (0) h |

\*Values in () are reported/estimated standard deviation.

**Supplementary Table S10:** Modified reduced-bioavailability for intraperitoneal IL-12 injection parameters fit to Lenzi et al. dataset.

| Parameter | Value |
| --- | --- |
| $k_{CB}(\text{days}^{-1})$ | 2.0 |
| $k_{CR}(\text{days}^{-1})$ | 0.46 |
| $k_{on}(\text{mL pmol}^{-1}\text{days}^{-1})$ | 5.4 |
| $k_{int}(\text{days}^{-1})$ | 0.46 |
| $N_L(\text{cells})$ | 1.0e9 |
| $\alpha(\text{pmol cells}^{-1}\text{days}^{-1})$ | 1.0e-5 |
| $R_0(\text{pmol cells}^{-1})$ | 1.7e-9 |
| $J_{LS}(\text{days}^{-1})$ | 0.94 |
| $J_{BL}(\text{days}^{-1})$ | 43 |
| $IC_{pstat}(\# \text{ cells}^{-1})$ | 9.1e2 |
| $\beta(\text{unitless})$ | 5.3e-3 |

**Supplementary Table S11:** Combined parameter set used to predict long-term, continuous dosing of IL-12 using local IL-12 production model.

| Parameter | Value |
| --- | --- |
| $k_{CB}(\text{days}^{-1})$ | 1.6 |
| $k_{CR}(\text{days}^{-1})$ | 0.46 |
| $k_{on}(\text{mL pmol}^{-1}\text{days}^{-1})$ | 200 |
| $k_{int}(\text{days}^{-1})$ | 0.56 |
| $N_L(\text{cells})$ | 3.8e10 |
| $\alpha(\text{pmol cells}^{-1}\text{days}^{-1})$ | 5.0e-6 |
| $R_0(\text{pmol cells}^{-1})$ | 5.6e-9 |
| $J_{LP}(\text{days}^{-1})$ | 0.94 |
| $J_{BL}(\text{days}^{-1})$ | 43 |
| $IC_{pstat}(\# \text{ cells}^{-1})$ | 1.4e6 |

**Supplemental Figure S1:**
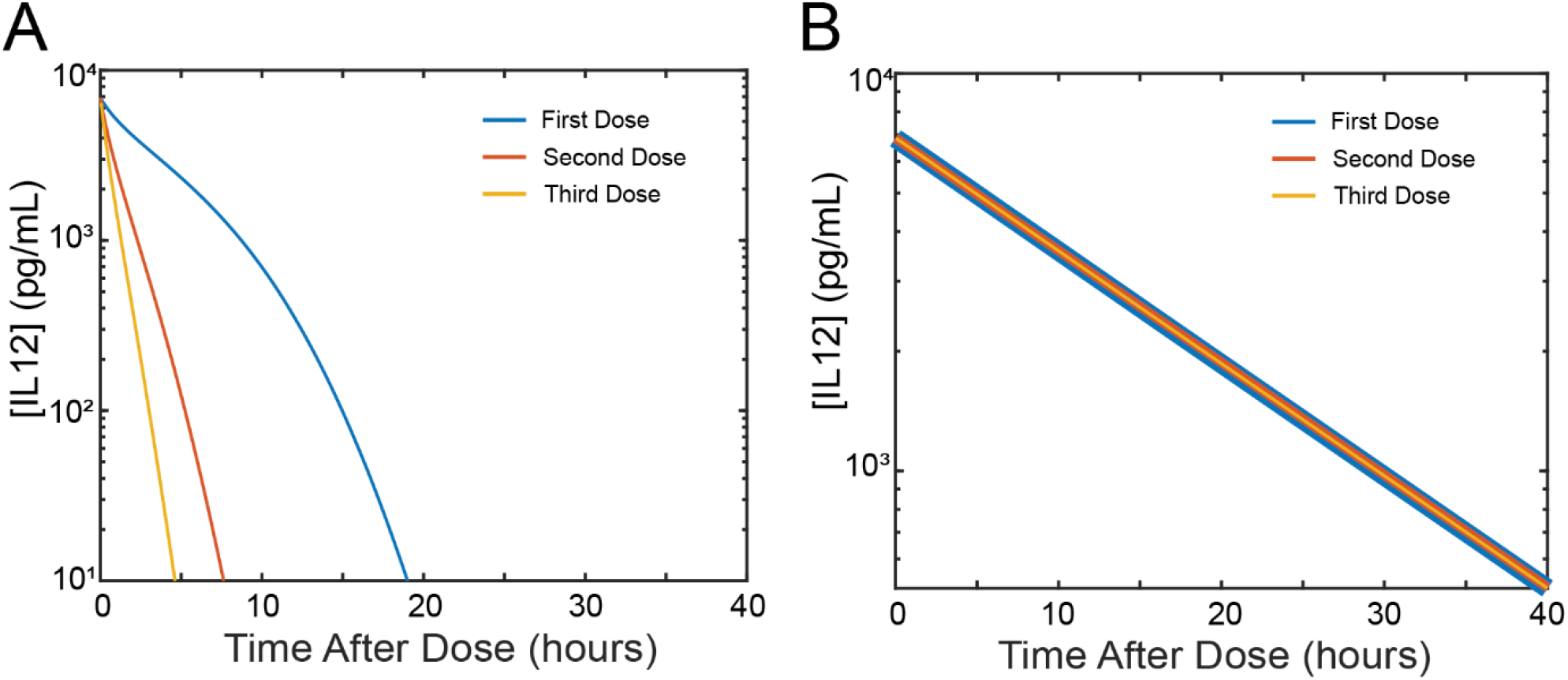
Simulation of repeated i.v. IL-12 dosing with model parameters fit to Motzer et al. dataset^1^. A) Predictions of repeated i.v. IL-12 dosing with the accelerated-clearance model. B) Predictions of repeated i.v. IL-12 dosing with the reduced-bioavailability model.

**Supplemental Figure S2:**
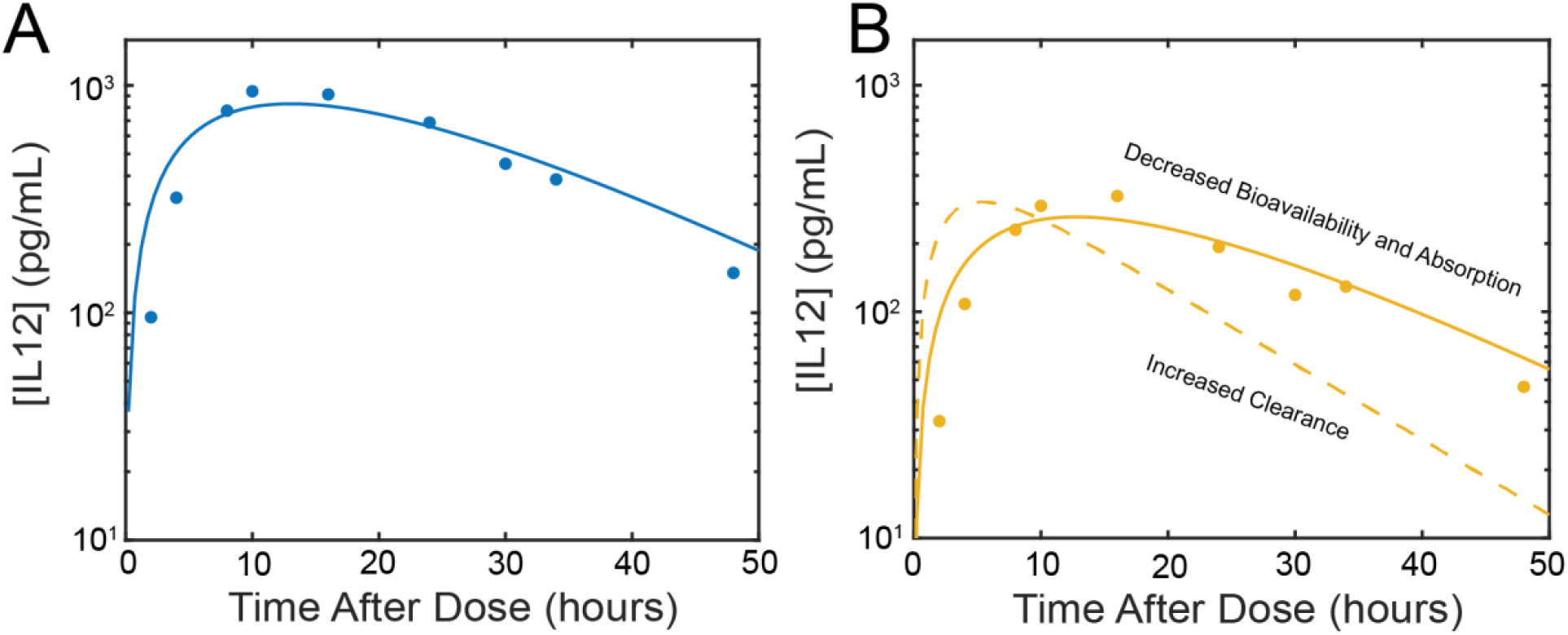
Modified accelerated clearance model without feedback fit to µg/kg datasets with and without prior exposure presented by Motzer et al. A) Model fit to dataset with no prior exposure estimating all parameters. B) Model fit to dataset with prior exposure varying only bioavailability and absorption (solid line) or clearance rate (dashed line) from the parameters in panel A).

**Supplemental Figure S3:**
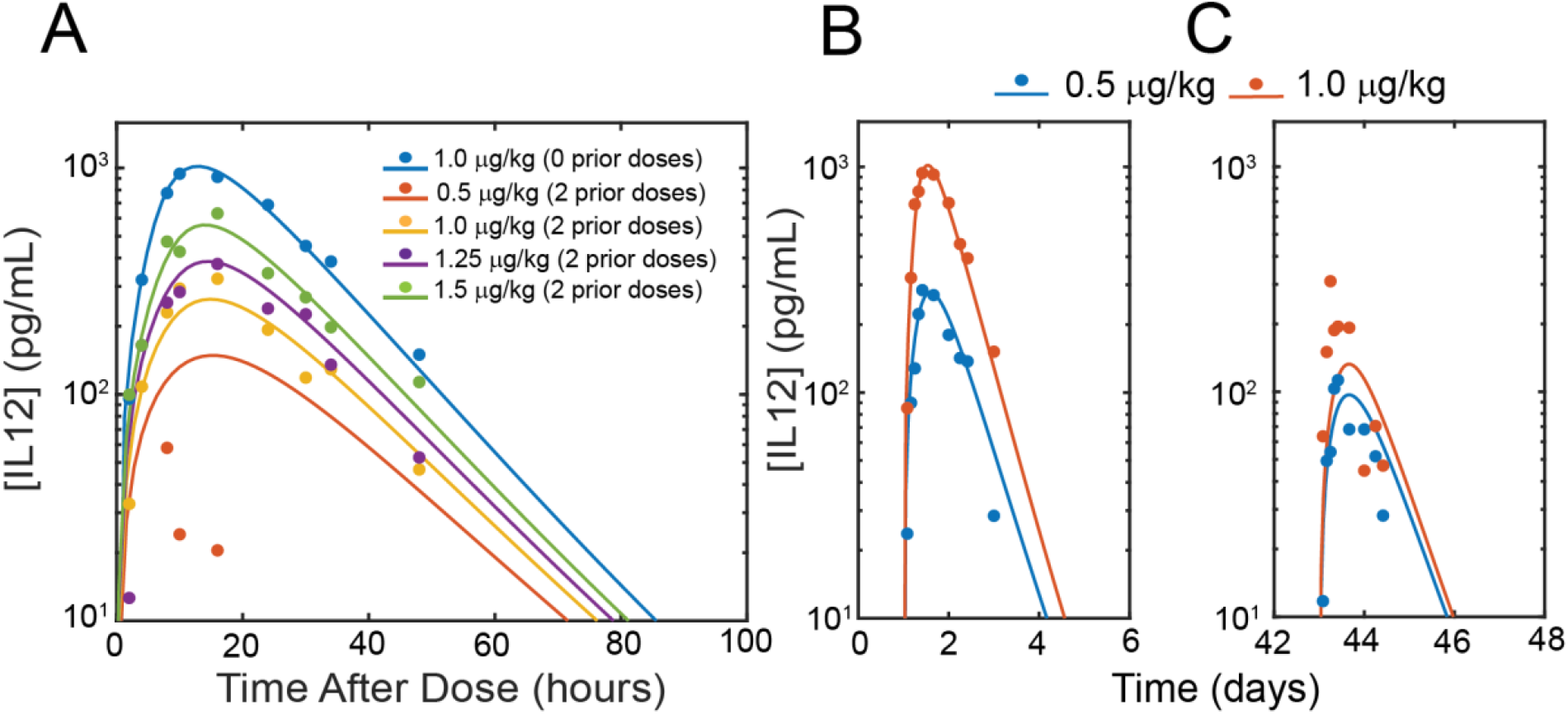
Reduced-bioavailability model fit to Motzer et al. and Rakhit et al. datasets simultaneously with global optimization. A) Model predictions compared to Motzer et al. data. B-C) Model predictions compared to Rakhit et al. data. B) Predictions and data after first dose. C) Predictions and data after sixth dose.

**Supplemental Figure S4:**
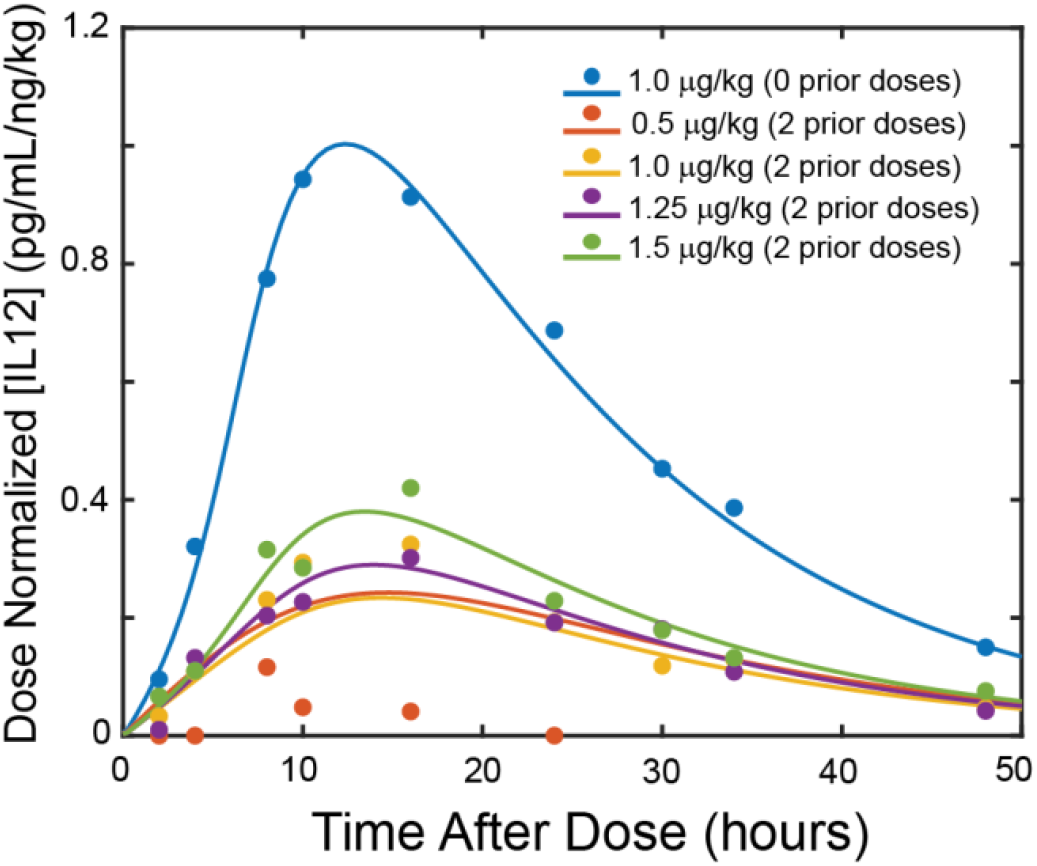
Reduced-bioavailability model predictions compared to clinical trial data presented by Motzer et al. with y-axis normalized by dose amount in ng/kg.

**Supplemental Figure S5:**
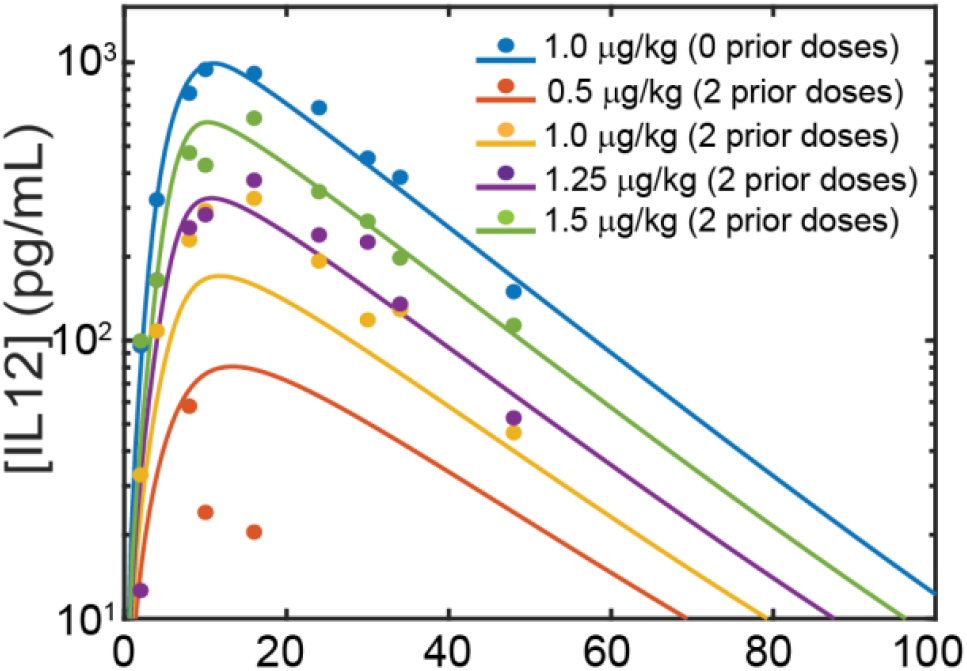
Addition of IL-12 receptor binding and upregulation in the blood compartment to the reduced-bioavailability model allows for capturing non-linearity in dose escalation.

**Supplemental Figure S6:**
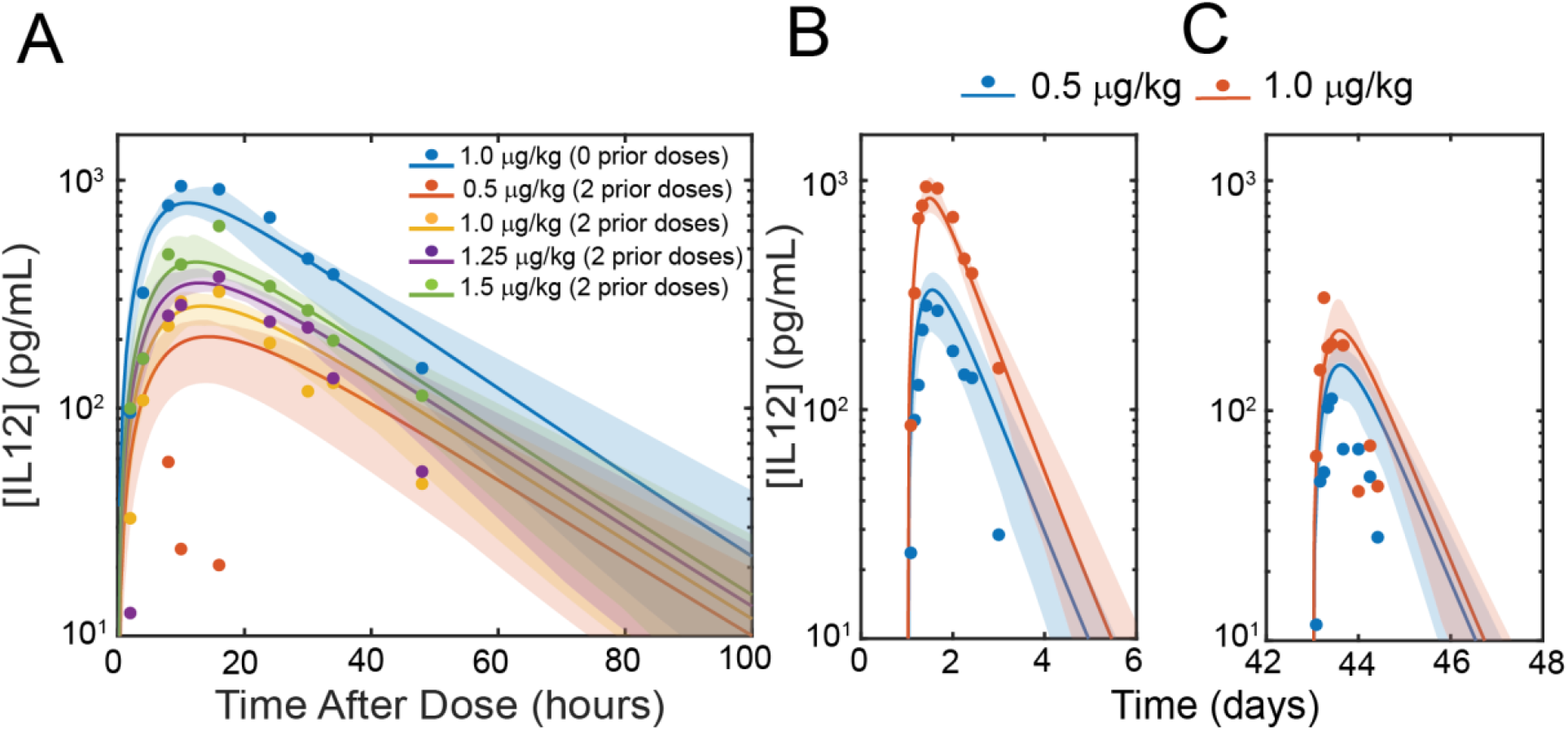
Reduced-bioavailability model fitness to clinical trial data following ensemble-based optimization. A) Model predictions fit to clinical trial data presented by Motzer et al.^1^. B-C) Model predictions fit to clinical trial data presented by Rakhit et al.^3^. Solid lines: mean predicted concentrations. Shaded area: raw 95% confidence interval.

**Supplemental Figure S7:**
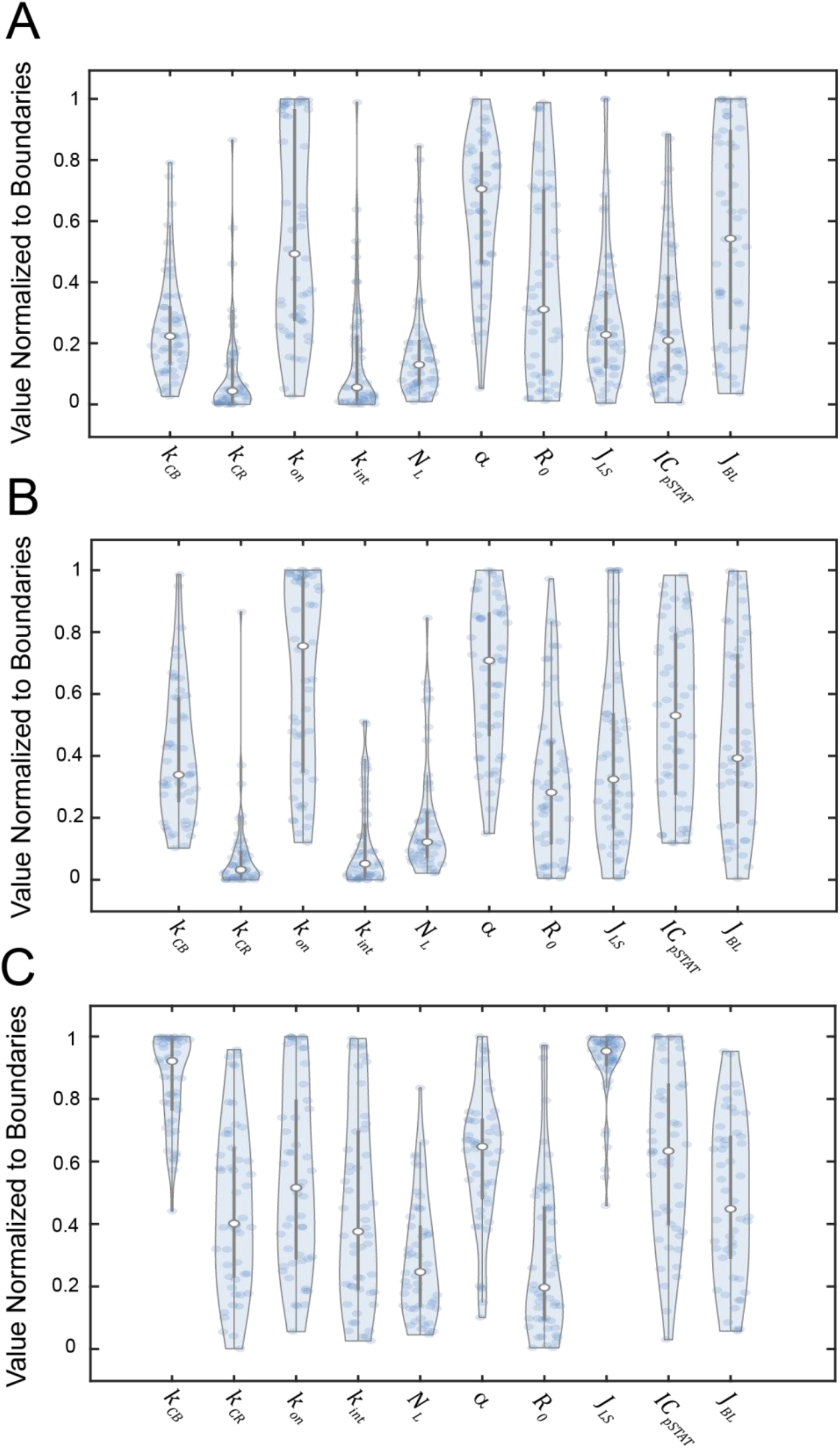
Distribution of locally optimized parameters fit to individual clinical trials. A) Distribution of 50 parameter sets locally optimized to Motzer et al.^1^ data. B) Distribution of 50 parameter sets locally optimized to Rakhit et al.^3^ data. C) Distribution of 50 parameter sets locally optimized to Portielje et al.^2^ data.

**Supplemental Figure S8:**
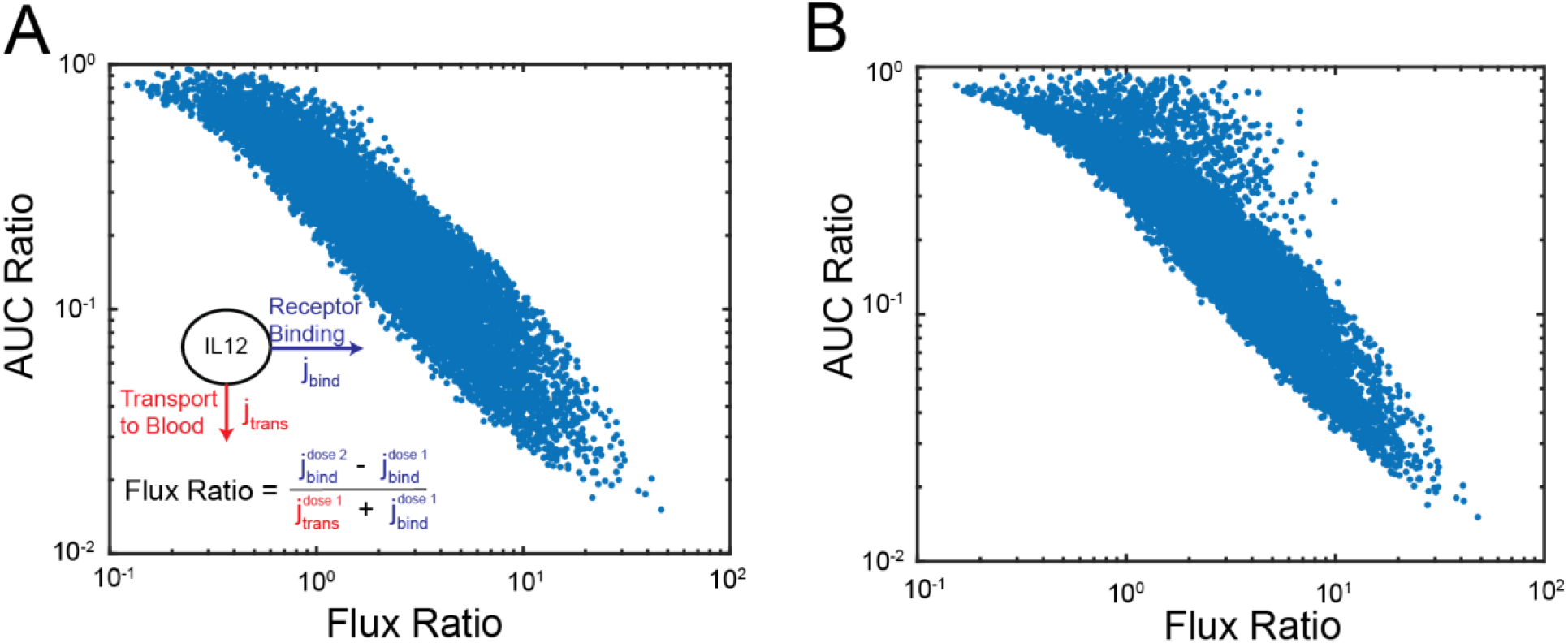
Relationship between binding flux ratio and amount of desensitization predicted by AUC ratio. A) Predicted amount of desensitization as quantified by AUC ratio v. flux ratio determined from numerical simulations. Inset: visual representation of flux ratio. B) Predicted amount of desensitization as quantified by AUC ratio v. flux ratio determined by analytical approximation of receptor dynamics (see SI Text).

**Supplemental Figure S9:**
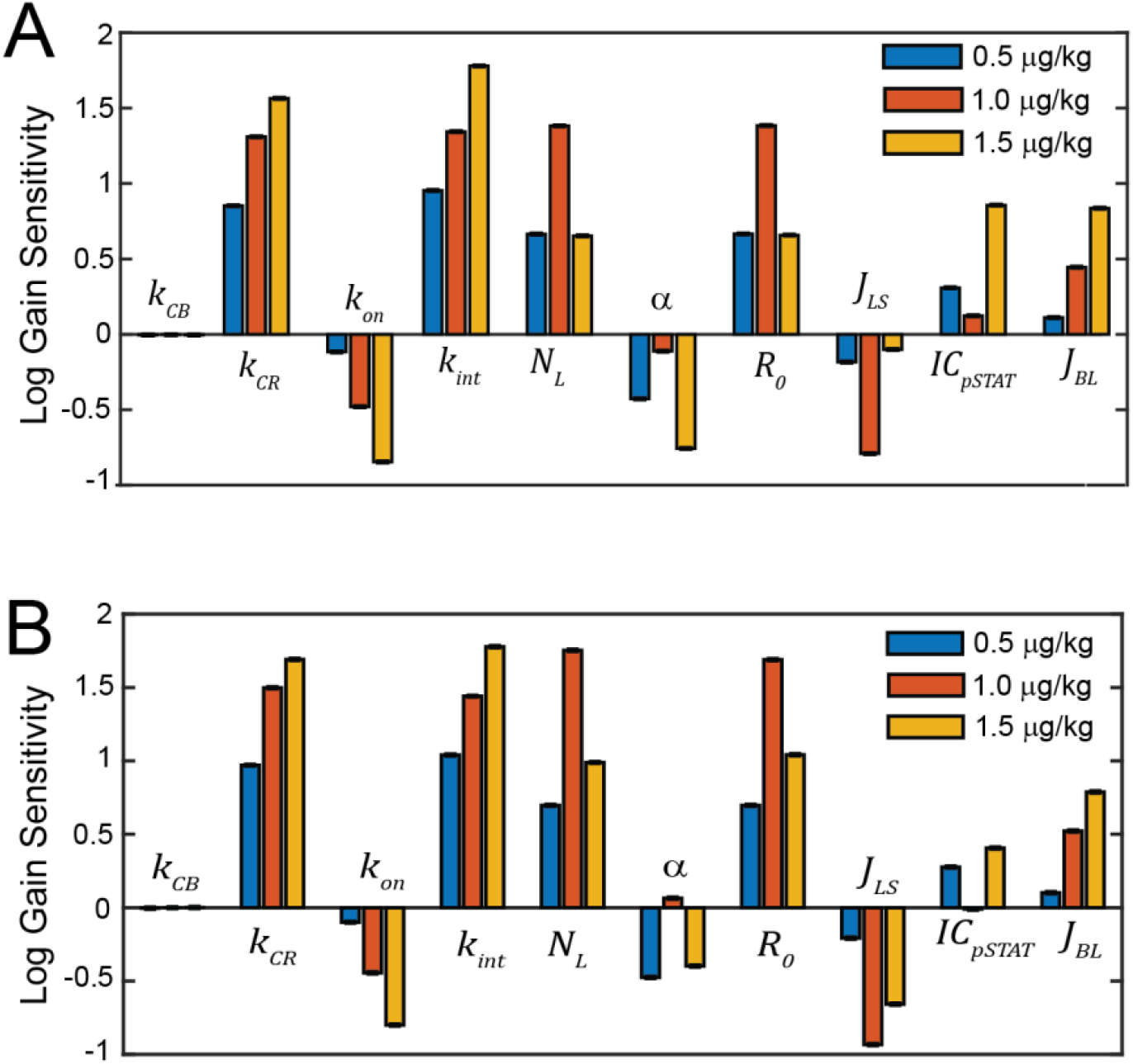
Amount of desensitization predicted by reduced-bioavailability model sensitivities to relevant model parameters. A) Desensitization log gain sensitivities after two consecutive 1.0 µg/kg doses. B) Desensitization log gain sensitivities comparing desensitization after three consecutive 1.0 µg/kg doses. Sensitivities are calculated according to equation S64.

**Supplemental Figure S10:**
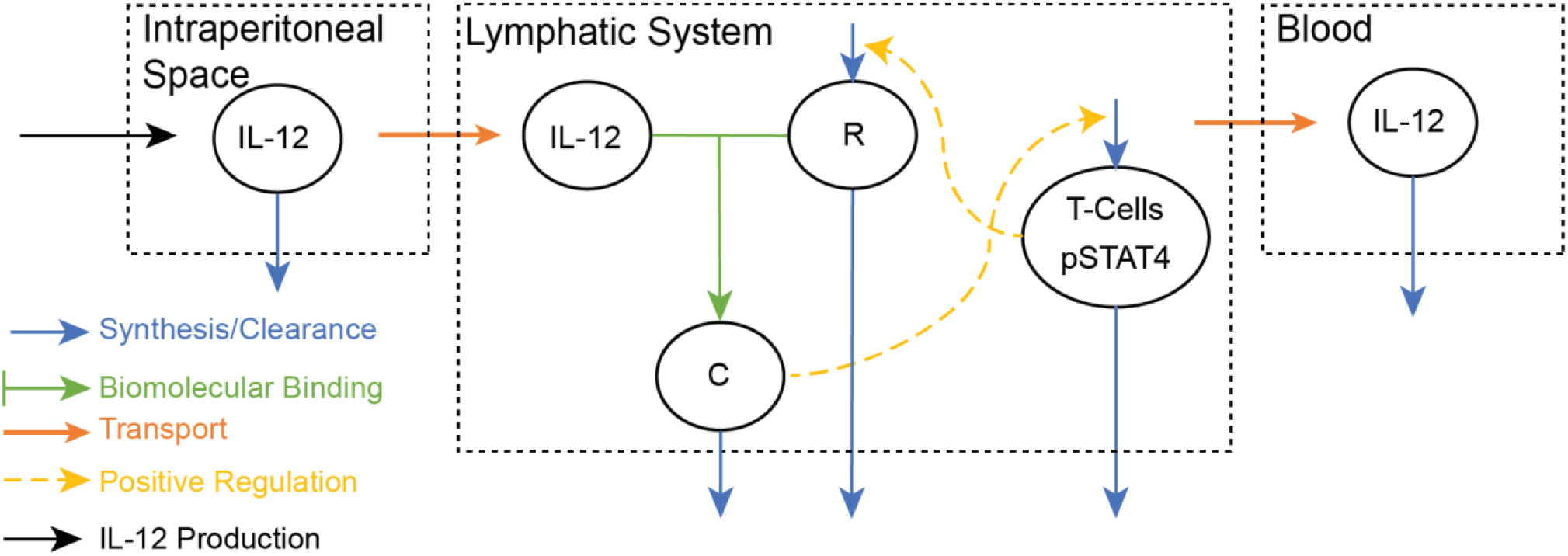
Schematic of local IL-12 production model. This model is identical to the reduced-bioavailability model with the dosing compartment volume, method of dosing, and transport parameters modified to represent continuous IL-12 production in the i.p. space.

**Supplemental Figure S11:**
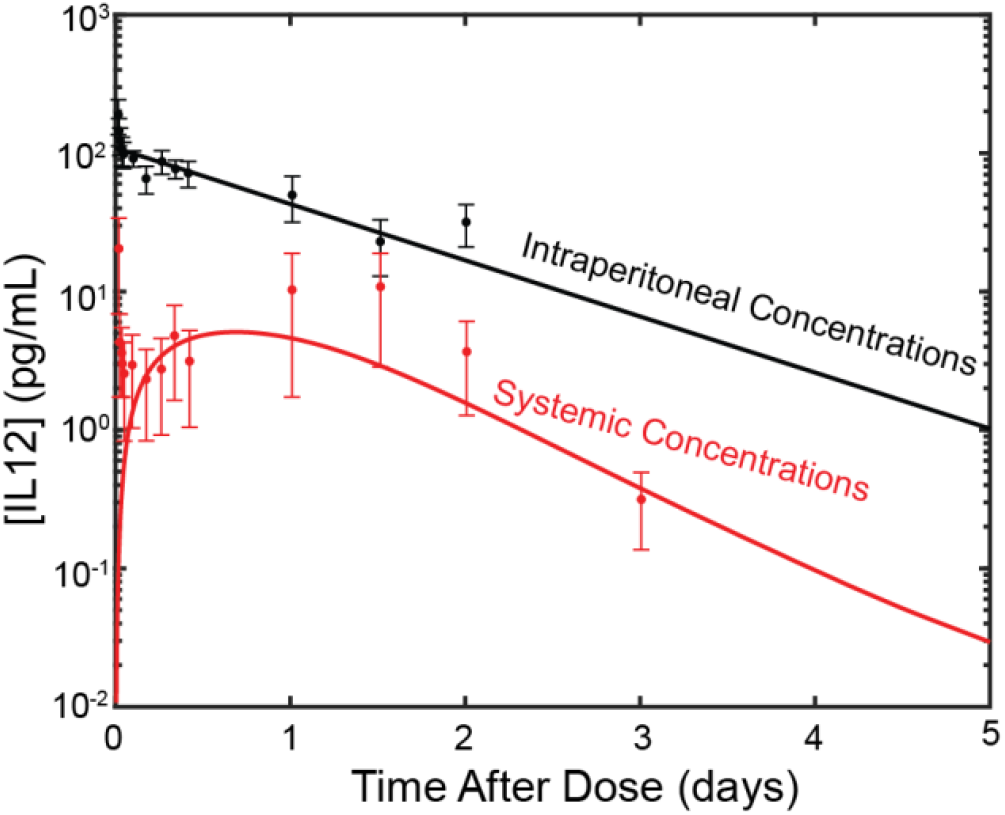
Reduced-bioavailability model modified to represent IP injection of IL-12 fit to clinical trial data.

**Supplemental Figure S12:**
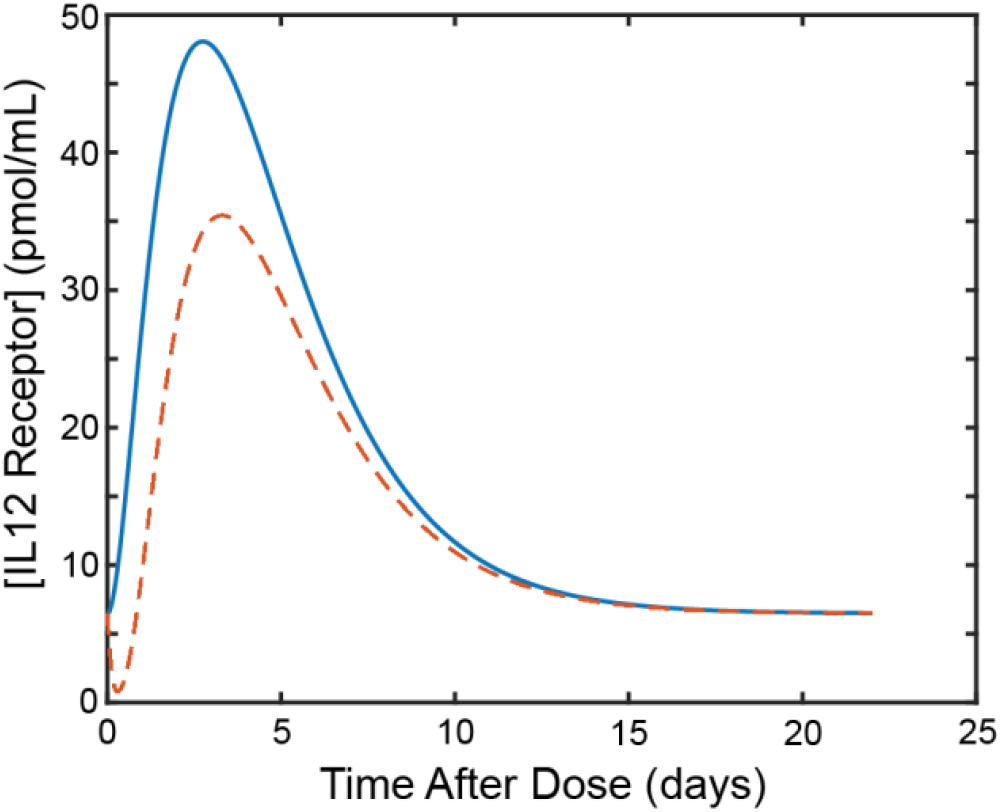
Analytical approximation for receptor concentrations compared to numerical simulations for reduced-bioavailability model. Solid blue curve: analytical approximation (see equations S74-S78). Dashed orange curve: numerical simulations.

**Supplemental Figure S13:**
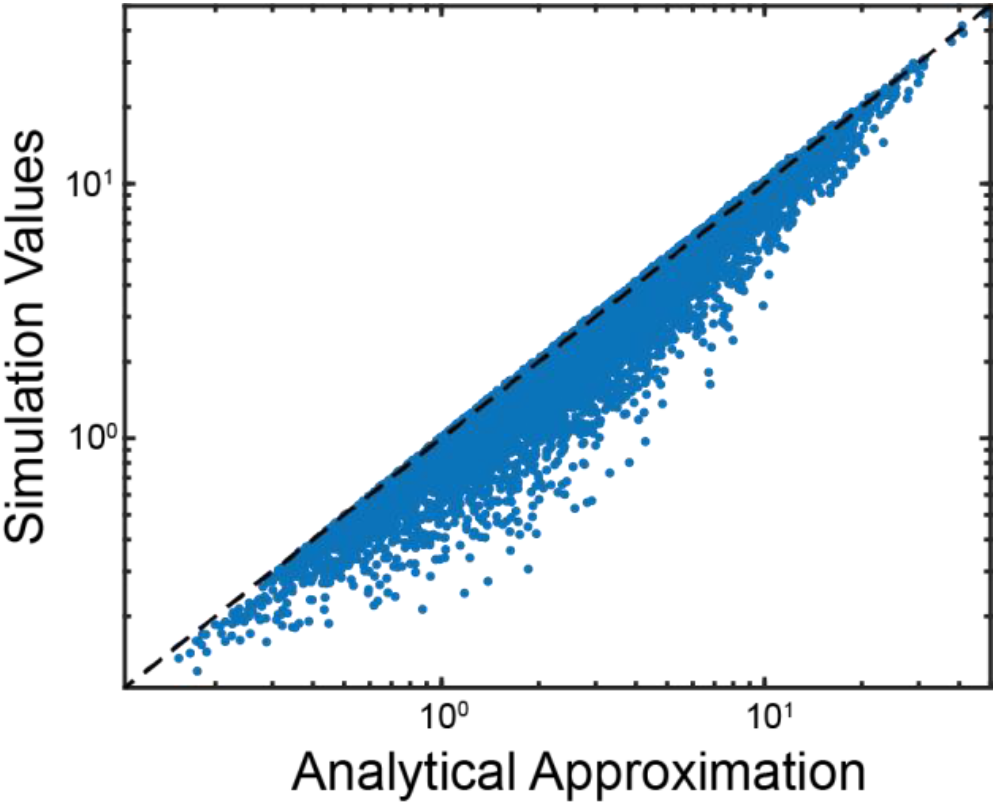
Comparison of determined IL-12 binding flux ratio (equations S65 and S66) as calculated by numerical simulation and our analytical approximation for receptor dynamics.

**Supplemental Figure S14:**
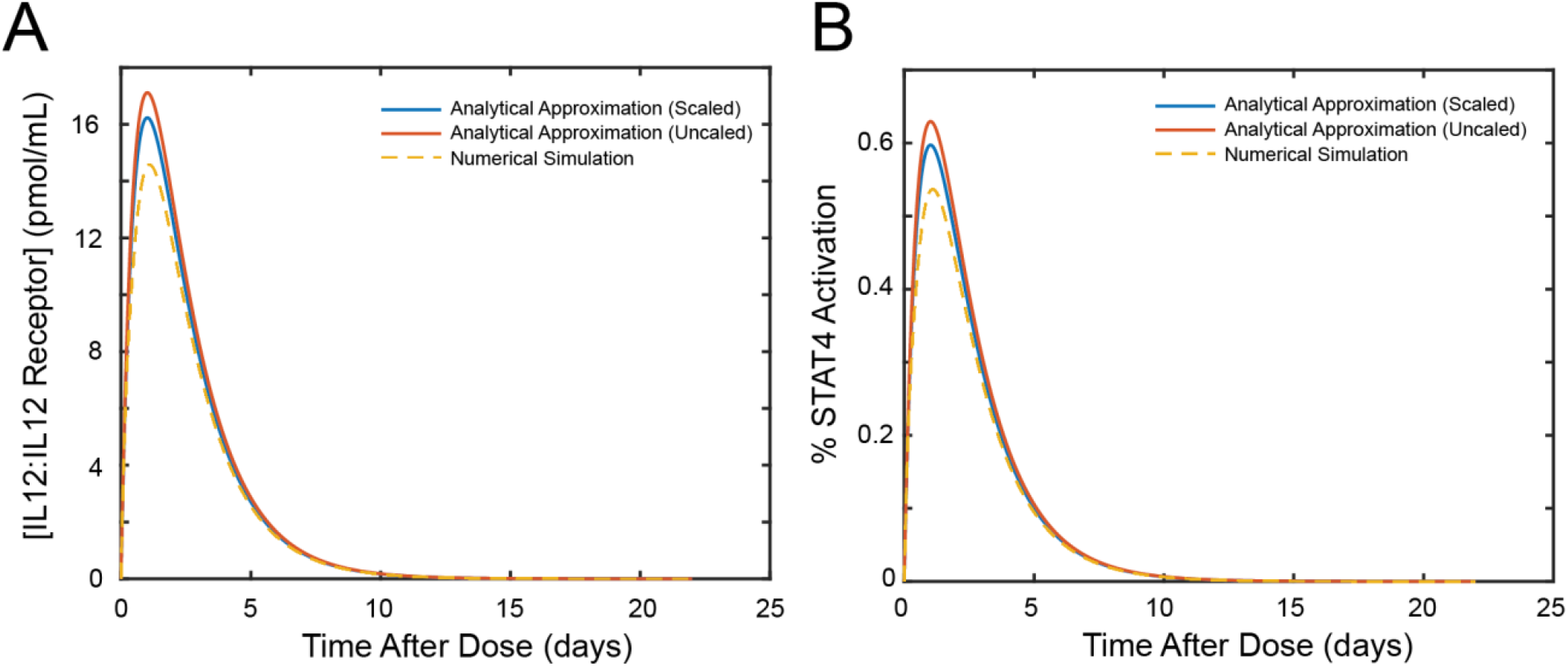
Comparison of analytical approximation for the concentration of IL-12:IL-12 receptor and STAT4 activation in the reduced-bioavailability model. A) Predictions of complex concentrations. B) Predictions of STAT4 Activation. Blue curves: Scaled analytical approximations. Orange Curves: Unscaled analytical approximations. Dashed yellow curves: Numerically simulated values. See equations S71-S72 for analytical expressions.

## List of Mathematical Models and Equations

*Accelerated-Clearance Model Equations*

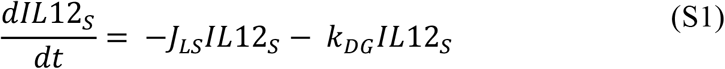

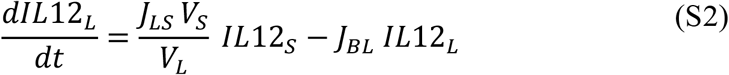

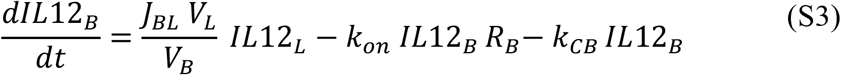

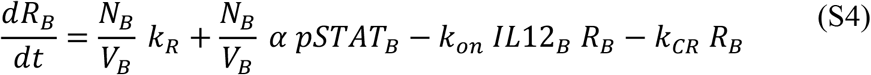

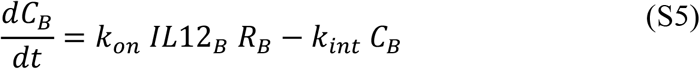

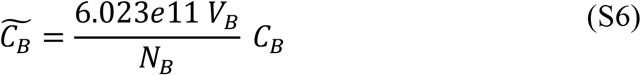

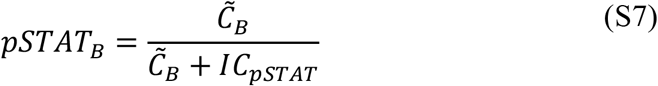

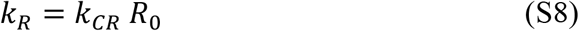

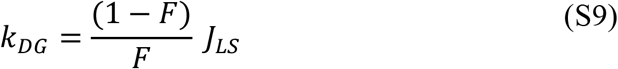

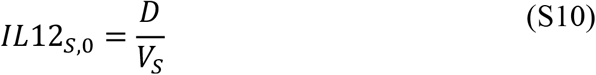

*Reduced-Bioavailability Model Equations*

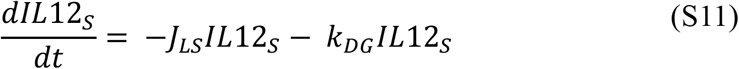

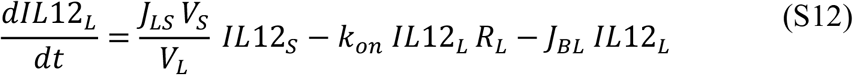

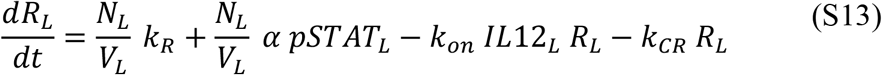

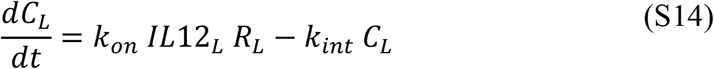

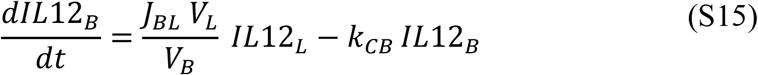

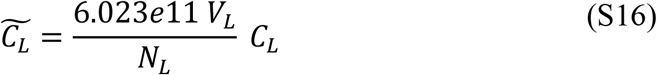

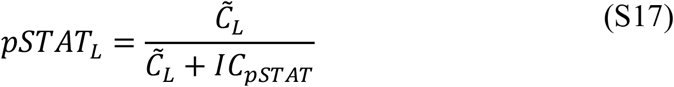

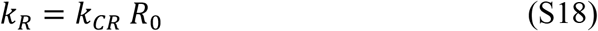

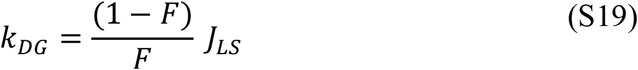

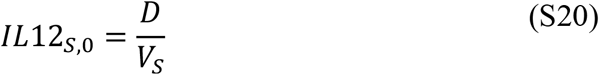

*Intraperitoneal Injection Model Equations*

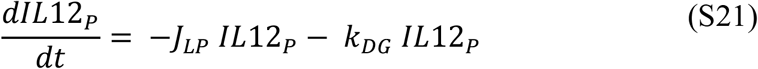

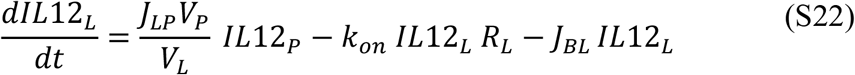

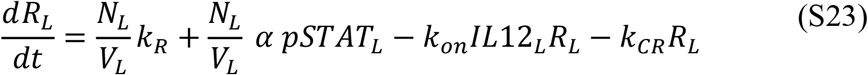

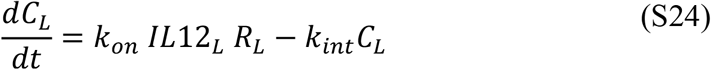

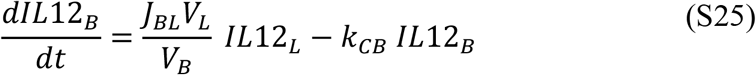

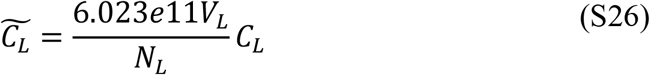

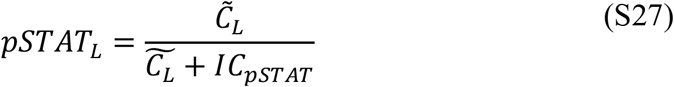

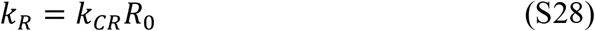

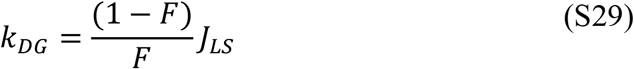

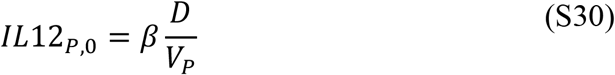

*Local Production Model Equations*

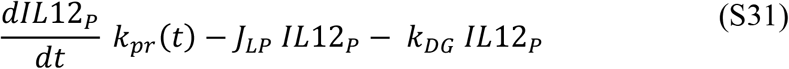

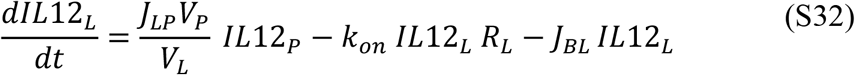

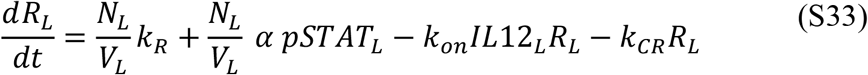

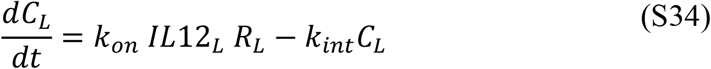

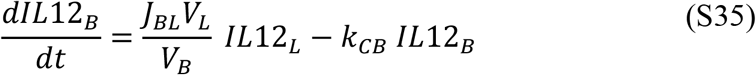

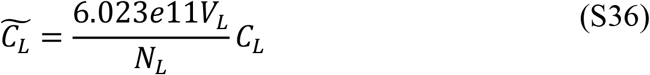

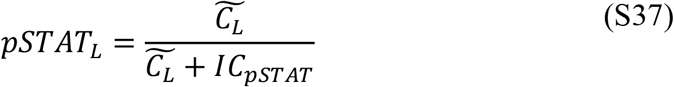

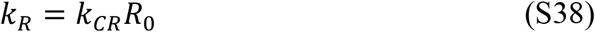

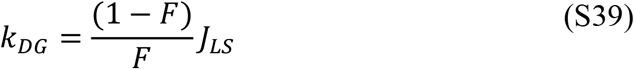

## Supplementary Text

### Derivation of Mathematical Models

The models used in this investigation are based on the same principles. Both the accelerated-clearance model and the reduced-bioavailability model consist of ordinary differential equations representing IL-12 transport via lymph flow from the subcutaneous space to the lymphatic system and to the blood and IL-12 receptor binding and upregulation. The key difference between the two models is the compartment where receptor binding occurs. The models used for the simulation of intraperitoneal IL-12 delivery are extensions of the reduced-bioavailability model. In the following sections, we derive the equations describing our models. The descriptions and values for the parameters used in each model are in **Supplementary Table S3**.

### IL-12 Dosing

IL-12 dosing occurs in the subcutaneous compartment. As the duration of injections is negligible compared to the simulation period, we model IL-12 dosing by setting the initial concentration of IL-12 in the subcutaneous space according to the following:

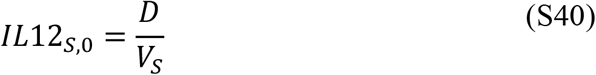

Here, *D* is the dose amount administered in pmol and *V*_*S*_ is the volume of the subcutaneous space in mL.

### IL-12 Transport Between Compartments

IL-12 transport between compartments is modeled based on a mass balance around each compartment and convective transport via lymph fluid flow. As the molecular weight of IL-12 is greater than 16 kDa, transport to the blood from the subcutaneous space primarily occurs through lymph flow^8^ and, therefore, is unidirectional. Consider a simple two-compartment system where lymph flow moves from compartment 1 to compartment 2 wit days^-1^. The mass balance around these two compartments is given by:

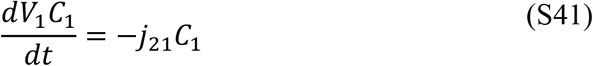

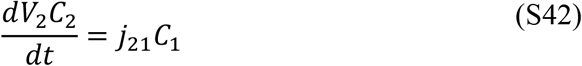

where *V*_*i*_is the volume of compartment *i* in mL and *C*_*i*_is the concentration of IL-12 in compartment *i* in pmol mL^-1^. We assume that the volume of each compartment remains constant, i.e., fluid moving from each compartment is instantly replaced or removed. Accordingly, we can rearrange the two differential equations to become:

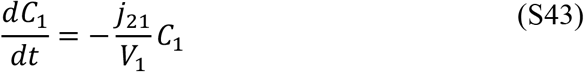

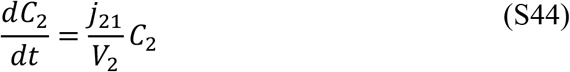

Making the substitution *j*_21_ = *J*_21_*V*_1_, we further simplify to:

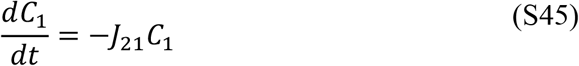

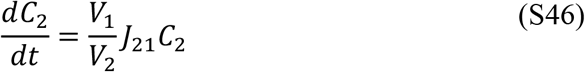

In these equations, *J*_21_ is the effective first order transport rate constant in days^-1^. Conceptually, this parameter refers to the fluid flow leaving compartment 1 normalized by the volume of compartment 1. This framework is generalized to represent transport between all compartments in our model.

### IL-12 Receptor Binding and Complex Formation

IL-12 binds its receptor either in the serum (accelerated-clearance model) or the lymphatic system (reduced-bioavailability model). For simplicity, we assume that binding is irreversible and occurs according to the rate law:

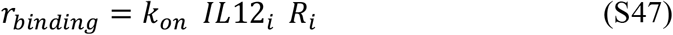

Here, *k*_*on*_ is the IL-12:IL-12 receptor association rate in mL pmol^-1^ days^-1^, *IL*12_*i*_is the concentration of IL-12 in compartment *i* where receptor binding occurs, and *R*_*i*_is the concentration of receptor in compartment *i* both in pmol mL ^-1^. In our model both IL-12 and IL-12 receptor are consumed according to *r*_*binding*_, and IL-12:IL-12 complex is generated at the same rate. Accordingly, the differential equation describing the concentration of IL-12:IL-12 receptor complex is given by:

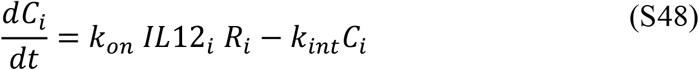

Here, *C*_*i*_is the concentration of IL-12:IL-12 receptor complex in compartment *i* and *k*_*int*_ is the first order complex internalization rate constant in days^-1^.

### STAT4 Activation

In both models, the formation of IL-12:IL-12 receptor complex leads to the activation of STAT4 based on the number of complexes per T-cell. Accordingly, we first calculate this quantity according to:

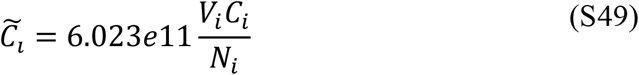

where 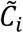 is the number of complexes per T-cell in # cells^-1^ in compartment *i, V*_*i*_is the volume of compartment *i* in mL, and *N*_*i*_is the number of T-cells in compartment *i*. The differential equation describing STAT4 activation is given by the following:

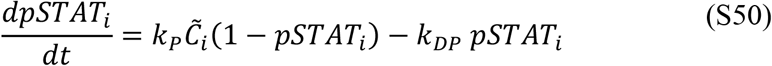

Here, *pSTAT*_*i*_is the fraction of phosphorylated (activated) STAT4, *k*_*p*_ is the first order phosphorylation rate in cell #^-1^ days^-1^, and *k*_*DP*_ is the first order dephosphorylation rate constant in days^-1^. Furthermore, as intracellular processes are much faster than physiological processes such as transport and clearance, we assume that STAT4 activation is in quasi-steady state in our models. Thus, we can solve for *pSTAT*_*i*_activation as the following:

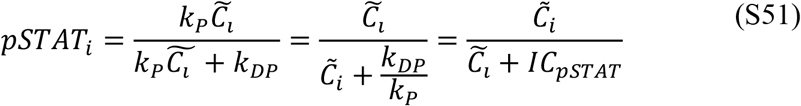

where *IC*_*pSTAT*_ is the number of complexes per T-cell required for half-maximum STAT4 activation and is equivalent to 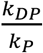.

### IL-12 Receptor Dynamics

In the compartment where receptor binding occurs, we represent the concentration of receptor based on a basal production rate, an upregulated production rate based on STAT4 activation, consumption via binding, and unbound internalization/clearance. Accordingly, the differential equation describing the concentration of receptor is given by:

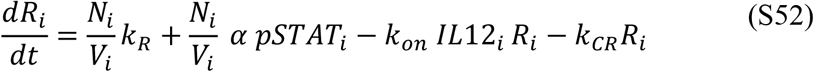

where *k*_*R*_ is the basal production rate per T-cell in pmol cells^-1^ days^-1^, *k*_*CR*_ is the unbound receptor internalization rate in days ^-1^, and *α* is the feedback parameter representing upregulated receptor production by STAT4 activation in pmol cells^-1^ days^-1^. Furthermore, we set the steady-state concentration of receptor according to the following relationship:

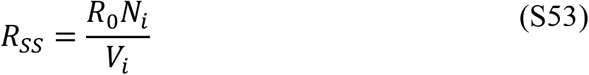

Here, *R*_*ss*_ is the steady-state/initial concentration of receptor in the absence of any IL-12 and *R*_0_ is the basal number of IL-12 receptors per T-cell in the absence of IL-12 and STAT4 activation. To ensure that the concentration of receptor remains at this level at steady state, we constrain the basal production rate *k*_*R*_ according to:

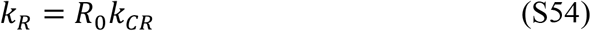

### IL-12 Clearance

In addition to receptor-mediated clearance, IL-12 can exit the body in our model via degradation in the subcutaneous space or traditional serum clearance. Clearance from the serum is determined by the first order clearance rate constant *k*_*CB*_ in days^-1^. Instead of directly specifying the degradation rate of IL-12 in the subcutaneous space, we specify the absolute bioavailability (fraction of IL-12 in the subcutaneous space that does not get degraded) and determine the corresponding degradation rate. Accordingly, the degradation of IL-12 in the subcutaneous space is determined by:

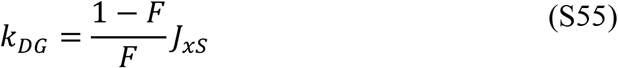

where *k*_*DG*_is the first-order subcutaneous degradation rate constant and *J*_*xS*_ is the first-order transport rate constant from the subcutaneous space to compartment *x*, both in days^-1^. In the accelerated-clearance model, this transport rate is that of IL-12 to the blood and in the reduced bioavailability model it is that of IL-12 to the lymphatic system.

### Model of Intraperitoneal IL-12 Injection

We developed the reduced-bioavailability model to simulate subcutaneous dosing of IL-12; thus, we modified our model slightly to fit intraperitoneal injection data presented by Lenzi et al.^9^. All model equations here are identical to the original model equations (other than slight notation changes) except the equation determining the initial concentration of IL-12 in the intraperitoneal cavity. In the clinical trial data presented by Lenzi and colleagues, the initial concentration of IL-12 in the intraperitoneal cavity is two orders of magnitude lower than expected given the reported dose amount. To account for this, we add the scaling factor *β* to our expression for the initial concentration of IL-12 in the intraperitoneal cavity to obtain:

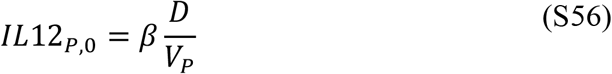

Here, *β* is a factor between zero and one that accounts for the discrepancy between the reported dose and initial concentrations of IL-12. Adding in this change and changes in notation to reflect IL-12 administration to the peritoneal cavity, we obtain equations S20-29

### Model of Continuous, Local IL-12 Delivery

We further modified our model to represent continuous, local delivery of IL-12 within the intraperitoneal cavity. All equations in this model are identical to that of intraperitoneal injection of IL-12 except for the equations governing IL-12 in the intraperitoneal space. Here, the ordinary differential equation describing the concentration of IL-12 in the intraperitoneal space is now given by:

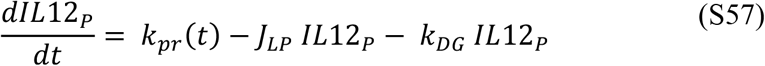

Here, *k*_*pr*_(*t*) is the 0^th^-order production rate of IL-12 that varies with time according to the dosing strategy being simulated in pmol days^-1^. We test three different dosing strategies designed to reach equivalent i.p. concentrations at long times: 1) constant production, 2) stepwise production, and 3) linearly increasing production. These production profiles are given by:

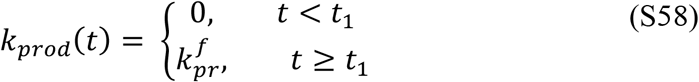

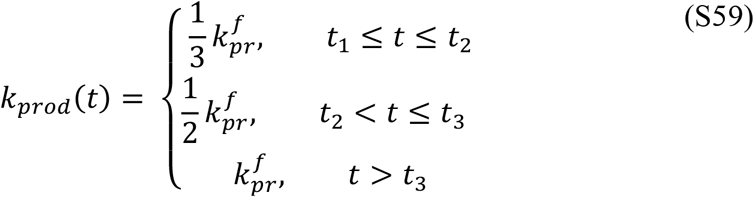

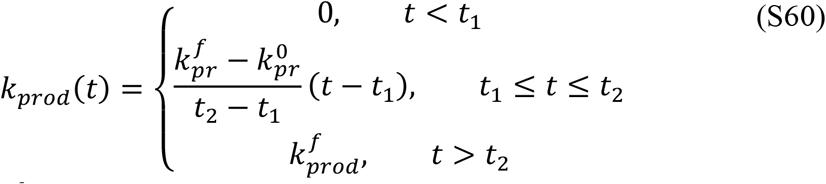

In the equations above 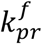 is the final production rate reached by each production profile, and *t*_1_ is the time dosing begins. The stepwise production profile is further characterized by *t*_2_ and *t*_3_, which are the times when IL-12 production is increased. Similarly, the linearly increasing production profile is characterized by *t*_1_ and *t*_2_, the time in which the increase in production rate begins and ends, respectively. All characteristic times of dosing profiles are given in days. Additionally, the initial production rate here is defined by 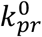.

### Parameter Optimization Error Functions

The error functions used for optimization depend on the type of data and the number of compartments for which data was available. For time course data presented for a single compartment, we used a modified version of the sum of standard squared error to account for the assay’s limit of detection:

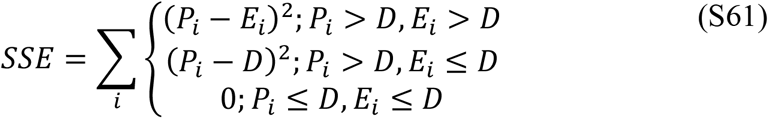

where *E*_*i*_is the value of experimental datapoint *i, P*_*i*_is the model predicted value of that datapoint, and *D* is the lowest limit of detection for the quantification assay. For time course data where data is presented for more than one compartment, i.e., blood and intraperitoneal IL-12 measurements, the error function was further modified to normalize to the magnitude of the experimental measurement according to:

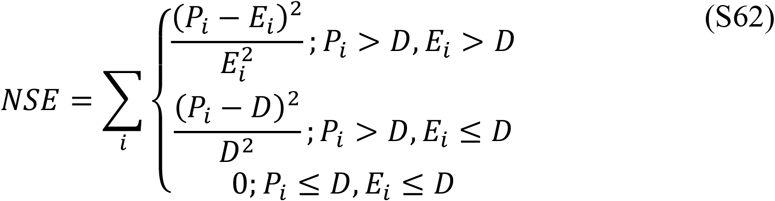

The error function used for optimizing to PK metric data is given by the following:

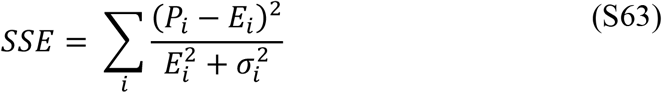

Here, *E*_*i*_is the value of the PK metric measured in the clinical trial, *σ*_*i*_is the reported standard deviation of the PK metric, and *P*_*i*_is the model predicted value.

### Local Model Sensitivity Analyses

Local sensitivity analyses were implemented to determine the impact of model parameters on the extent of IL-12 PK desensitization. For the simulation of multiple repeated doses of the same amount, we quantify the amount of desensitization predicted as the ratio of IL-12 serum AUC after the last and first dose according to:

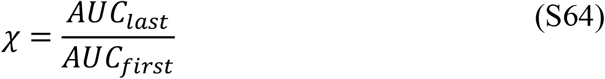

To determine the sensitivity of the quantity *χ* to a given parameter *p*, we calculated the log gain sensitivity according to:

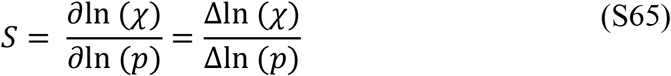

### Ensemble Analysis of Desensitization Requirements

To expand on the influence of the parameters related to receptor lifetime, initial receptor concentrations, and feedback dynamics we generated an ensemble of 10,000 parameter sets varying *k*_*CR*_, *k*_*int*_, *N*_*cells*_, *α, R*_0_, and *IC*_*pSTAT*_ between 50% and 150% of the optimized values holding all other parameters constant. Each parameter set was simulated for two consecutive 1.0 µg/kg doses on days 1 and 8 of a 15-day cycle, and the amount of desensitization predicted was quantified based on equation 24. Analysis of these simulations illustrated that the amount of desensitization that occurs is correlated to the increase in binding flux at the time of the second dose (t = τ) normalized by the total flux of IL-12 out of the lymphatic system at the time of the first dose (t = 0), determined by equation 30. More specifically, we found that the value Ψ was inversely correlated to the AUC ratio and, therefore, positively correlated with the extent of desensitization predicted (**Supplemental Figure S8A**)

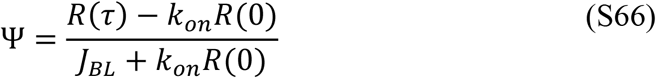

To gain deeper insight into the role specific parameters have on the amount of receptor remaining at the time of the second dose and the amount of desensitization predicted, we derived an analytical approximation for the concentration of receptor in the lymphatic system following a single dose (equations S74-78) This approximation of receptor concentrations is in reasonable agreement with numerical simulations, especially at longer time points where the receptor concentration is close to baseline levels (**Supplemental Figure S12**). Using this approximation to calculate the increase in receptor binding flux at the time of second dose based on equation S65, we obtain:

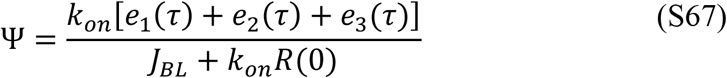

Here, the three exponential terms *e*_1_(*t*), *e*_2_(*t*), and *e*_3_(*t*) are given by equations S75-77. When used to predict the magnitude of IL-12 desensitization that occurs, equation S66 yields similar results to those extracted from numerical simulations (**Supplemental Figures S8B and S13**). Our analytical expression illustrates how specific parameters influence the amount of desensitization predicted without numerical simulation. We see that terms in the numerator of *ϕ* will lead to a larger peak in receptor concentration, and vice versa for the denominator. Additionally, increases in the three exponential decay parameters, *J*_*LS*_, *k*_*int*_, and *k*_*CR*_, lead to faster receptor clearance. If all three decay timescales are sufficiently short, upregulated receptor will clear by the time of the second dose and desensitization will not be predicted. Furthermore, we predict that multiple processes in our model contribute to the presence of desensitization, and that desensitization will be predicted over a variety of different parameter combinations. Generally, the requirement for desensitization in our model is that the upregulated of IL-12 receptor remains present when the next dose is administered. We show here that this behavior is not specific to a small subset of parameter values, supporting the robustness of our model to predict desensitization despite the inherent variability in our model parameters.

### Derivation of Analytical Approximation for Receptor Dynamics

To approximate the concentration of IL-12 receptor in the reduced-bioavailability model over time with an analytical expression, we aimed to approximate species in the model with exponential expressions. As the upregulation of receptor is regulated by the amount of STAT4 activation in T-cells, which is in turn regulated by the amount of IL-12:IL-12 receptor, we start by approximating the amount of complex formation that occurs in the lymphatic system. The ordinary differential equation describing the concentration of complex formed is given in equation S13. Importantly, this differential equation is nonlinear due to the *k*_*on*_*IL*12_*L*_*R*_*L*_ term. As a linear approximation, we assume that the binding flux is equal to the flux into the lymphatic system such that *k IL*12 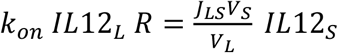, giving us:

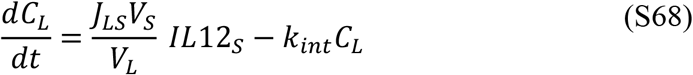

The concentration of IL-12 in the subcutaneous space, *IL*12_*S*_, is described by a single exponential decay since for this model we assume bioavailability (*F*) is one. The exponential decay describing the concentration of IL-12 in the subcutaneous space is given by:

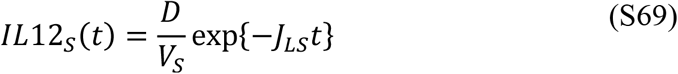

Making this substitution allows us to solve equation S67 analytically to obtain the dual exponential decay given by:

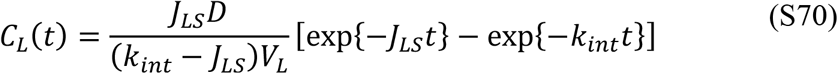

Based on equations S15 and S16, we can now approximate the level of STAT4 activation according to:

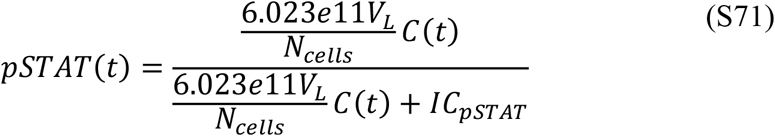

These approximations for the concentration of complex and STAT4 activation are in reasonable agreement with numerical simulations (**Supplemental Figure S14**). However, there is a slight overprediction of both species, likely due to our assumption that binding flux is equal to the total flux of IL-12 into the lymphatic system. To account for this difference, we rescale our approximation for the amount of complex formed by the initial ratio of binding flux to total flux, such that *C*_*L*_(*t*) is now given by:

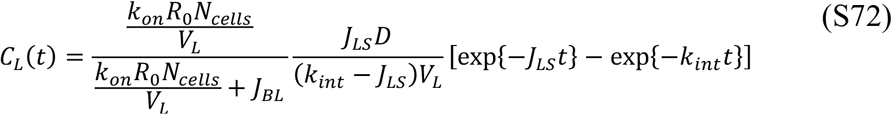

As seen in **Supplemental Figure S14** (blue curves), this slightly increases the accuracy of our analytical approximations compared to numerical simulations. Using equations S13 and S71, we can now solve for an analytical approximation for the concentration of receptor in the lymphatic system after linearizing the equation. Here, we make an important assumption that at long times (around the time of the next dose, which is our time span of interest), the amount of IL-12 in the lymphatic system is negligible such that *k*_*on*_*IL*12_*L*_*R* ≈ 0. Additionally, our expression for *pSTAT*(*t*) is non-linear; however, in the parameter regime we are operating 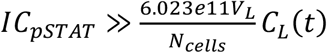, thus we can assume this function is approximately linear as given by:

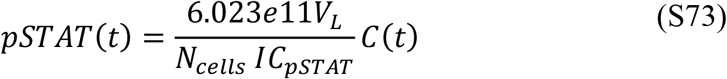

Thus, our ordinary differential equation for receptor concentrations in the lymphatic system becomes linear, given by:

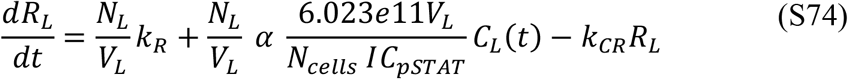

This ordinary differential equation can be solved analytically and making the important substitution *k*_*R*_ = *k*_*CR*_*R*_0_, we obtain:

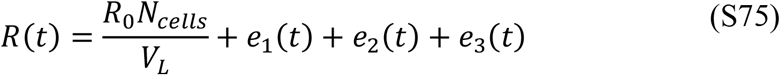

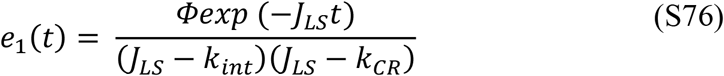

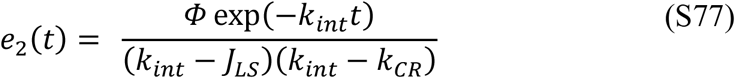

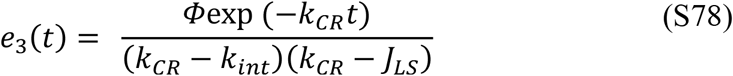

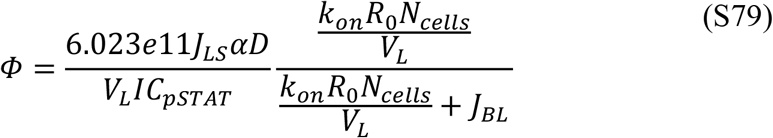

## Acknowledgments

An early version of this work was presented as a late-breaking abstract at the BMES 2023 Annual Meeting.

This work was partly supported by the Welch Foundation (grant C-1995 to O.A.I.) and by the Advanced Research Projects Agency for Health (ARPA-H) under Award Number AY1AX000003. The simulations were performed on the Rice University’s Center for Research Computing equipment acquired with the Big-Data Private-Cloud Research Cyberinfrastructure MRI-award funded by the NSF under grant CNS-1338099. The content is solely the responsibility of the authors and does not necessarily represent the official views of the Welch Foundation, the Advanced Research Projects Agency for Health, or the NSF.

## Notes

**Conflicts of Interest** O.V. holds equity and O.A.I has consultant position in Avenge Bio. The authors declare that they have no other competing interests.

### Competing Interest Statement

O.V. holds equity and O.A.I has consultant position in Avenge Bio. The authors declare that they have no other competing interests.

### Summary of Updates

We have revised our discussion and development of the accelerated-clearance model

